# Neutrophils reprograms the bone marrow to impair T-cell immunity during tuberculosis

**DOI:** 10.1101/2022.06.28.498035

**Authors:** Mohd. Saqib, Elizabeth McDonough, Shreya Das, Zhongshan Cheng, Poornima Sankar, Fuxiang Li, Qianting Yang, Yuzhong Xu, Weifei Wang, Xinchun Chen, Anil K Ojha, Fiona Ginty, Yi Cai, Bibhuti B Mishra

## Abstract

*Mycobacterium tuberculosis* (Mtb) infection induces persistent influx of neutrophils that associates with poor bacterial control and clinical outcome from tuberculosis (TB). Although implicated in TB pathology, the mechanism by which these cells contribute to pathogenesis is poorly understood. Using Cell-DIVE multiplexed immunofluorescence imaging and spatial analysis of inflammatory TB lesions, we demonstrated that persistent neutrophil infiltration affects the spatiotemporal organization of T-lymphocytes and impairs their function. Instead of directly suppressing T-cells, neutrophils produce granulocyte colony stimulating factor (CSF3/G-CSF) that collaborates with type I interferon (IFN-I) to promote a granulocyte-skewed hematopoiesis impacting T-lymphocyte production. Importantly, neutrophil-intrinsic IFN-I receptor 1 (IFNAR1) is both necessary and sufficient to promote pathologic granulopoiesis. Finally, inhibition of IFNAR1-signaling alone mitigates immunopathogenesis by restoring hematopoietic equilibrium. Collectively, our work uncovers a potential immunevasion strategy by which virulent Mtb strains induce IFN-I to generate pathogen-permissive neutrophils that produce G-CSF and sustain pathogenic hematopoiesis to impair T-cell immunity during TB.

---

To survive a chronic infection, the host must optimally control pathogen replication and limit the collateral tissue damage caused by overwhelming inflammatory response. Deregulation of such protective immune responses to chronic *Mycobacterium tuberculosis* (Mtb) infection often associates with immunopathology resulting in the inflammatory disease tuberculosis (TB)^1,2^. Identification of the factors that tip the balance between a protective and pathological immune response during TB is crucial for designing interventions and to predict potential TB-associated complications of immune therapies used against other human diseases.

While specific causal factor(s) regulating TB pathogenesis are currently unknown and likely multifarious, emerging evidence from studies in humans and experimental animal models have underscored the importance of interferons (type I and II) and interleukin-1 (IL1) for host defense against TB^3^. Defects in IFNγ production or signaling causes susceptibility to Mtb infection^4–7^. The protective effects of IFNγ is partly due to its ability to induce nitric oxide synthase 2 (iNos) that curtails IL-1 dependent inflammatory tissue damage. However, mice deficient in IL-1 signaling succumb to severe TB that has been attributed to failed bacterial control and unrestrained inflammation mediated by deranged eicosanoid signaling^8–10^ suggesting that the protective and pathological function of IL-1 is dependent on the stage of infection, and magnitude of the signaling. In contrast, the type I IFN (IFN-I) response is a marker of disease progression as it counter-regulates the protective immunity conferred by both IFNγ and IL1^10,11^. Mutations in the ligand binding domain of the IFN-I receptor 1 (IFNAR1) that dampens the signaling are associated with protection from active TB whereas hypermorphic allele that contributes to a stronger signaling response enhances the risk of pulmonary TB in humans^12,13^. Susceptibility due to dysregulation of the IFN-I/IL1 axis is observed in C3HeB/FeJ mice, a strain that develops human like TB lesions^14^. The pathological IFN-I response is tightly controlled by Sp140, a transcription factor encoded by the super susceptibility to tuberculosis 1^R^ (sst1^R^) haplotype in the resistant C57BL/6 mice compared to the C3HeB/FeJ mice^15^. One of the IFN-I inducible genes that is expressed in these mice and phenocopies the sst1^s^ locus with regard to TB susceptibility is the IL-1 receptor antagonist (IL-1R1n) that impairs an optimal IL1 immunity^16^. Thus, the intricate balance between protective (IL-1 and IFNγ) and pathological (IFN-I) immune responses determine TB disease susceptibility in mice and humans.

A common feature of severe TB in both humans^17,18^ and mice with alteration in IFNs and IL-1 levels is the infiltration of neutrophils to the lung^7,19,20^. While neutrophil recruitment is critical for protective immunity to many pulmonary pathogens, multiple lines of evidence in humans, non-human primates (NHPs), and mice indicate that these cells are pathologic during TB^21,22^. Depletion of neutrophils from a variety of susceptible mouse strains reduces immunopathology and restores host control of bacterial replication. Neutrophils represent the predominant infected cells in tissue biopsies of TB patients^23^ and most notably, gene signatures related to neutrophil function are strong predictor of disease progression and poor prognosis^17,24^. While unchecked neutrophil influx is regarded as detrimental, the abundance and spatial arrangement of T and B lymphocytes in TB lesions has a positive impact on the anti-microbial immunity^25^. Mice lacking T- and B-cells, TCRα/β signaling, MHC-II are highly susceptible to TB^26^. Both mouse and NHP lungs containing lymphoid follicles called inducible bronchus associated tissue (iBALT) are associated with better bacterial containment^27^. Multimodal profiling of lung granulomas in cynomolgus macaques identified Th1-Th17, stem like T-cells as cellular corelates of bacterial control^28^. Altogether, these observations advocate for a model where T-cell responses in TB lesions are protective and neutrophil-dominant inflammatory responses are detrimental. However, the underlying regulatory mechanism of this protective and pathological immune responses induced by Mtb infection is poorly delineated.

Here, we have shown that Mtb HN878 infection induces IFNAR1-dependent granulocyte-biased hematopoiesis that negatively impacted the lymphocyte production, and ultimately the T-cell immunity in the lung and spleen. We have further demonstrated that neutrophil-intrinsic IFN-I signaling is both necessary and sufficient for governing the infection-induced granulopoiesis and recruitment of neutrophils to the lung. Our work highlighted for the first time the role of neutrophils as bonafide producers of G-CSF that sustains the pathologic granulopoiesis mediated by IFN-I. Thus, our work has uncovered a novel immunevasion strategy by which Mtb induces IFN-I to generate pathogen permissive neutrophils and by a G-CSF dependent feed-forward mechanism sustains the pathologic granulopoiesis to impair protective T-cell immunity during TB.

## Results

### Higher neutrophil to T-lymphocyte ratio accurately predicts disease outcome in mice

To dissect the host determinant(s) of protective and pathological immune responses during TB, we evaluated the disease in mice following low dose (~100 CFU) aerosol challenge of Mtb strain HN878 that belongs to the lineage 2 of the W/Beijing isolates^29^. Mtb HN878 infection induces highly inflammatory TB in the relatively resistant C57BL/6 mice (referred as BL/6) compared to the commonly used laboratory strains^30,31^. Infection with Mtb HN878 causes severe disease in C3HeB, *iNos^−/−^* and *Il1r1^−/−^* mice (**Extended Data Fig.1a**). All these animals rapidly succumb to TB due to higher bacterial burden, neutrophil influx, and significantly fewer T-cells (CD3e+) in the lungs compared to BL/6 mice. The inverse trend in neutrophil and T-cell numbers was reflected in a higher neutrophil to T-lymphocyte ratio (referred to as NLR) that correlates with the susceptibility of mice (**Extended Data Fig. 1b-e**).

### Abundance of neutrophils impairs T-cell response in TB lesions

Since high NLR was a strong predictor of disease outcome in mice, we sought to determine whether the NLR has any relevance to disease status in human pulmonary TB. High neutrophil, low lymphocyte count reflected in a greater NLR was observed in the peripheral blood of TB patients compared to healthy cohorts. Importantly, the neutrophil counts and NLR were positively correlated with the severity of lung disease measured by sputum smear AFB positivity, HRCT scores, and ESR in TB patients (**Fig. 1a-c, Extended Data Fig. 1f-h**), consistent with the observations in mice.

**Fig. 1.**
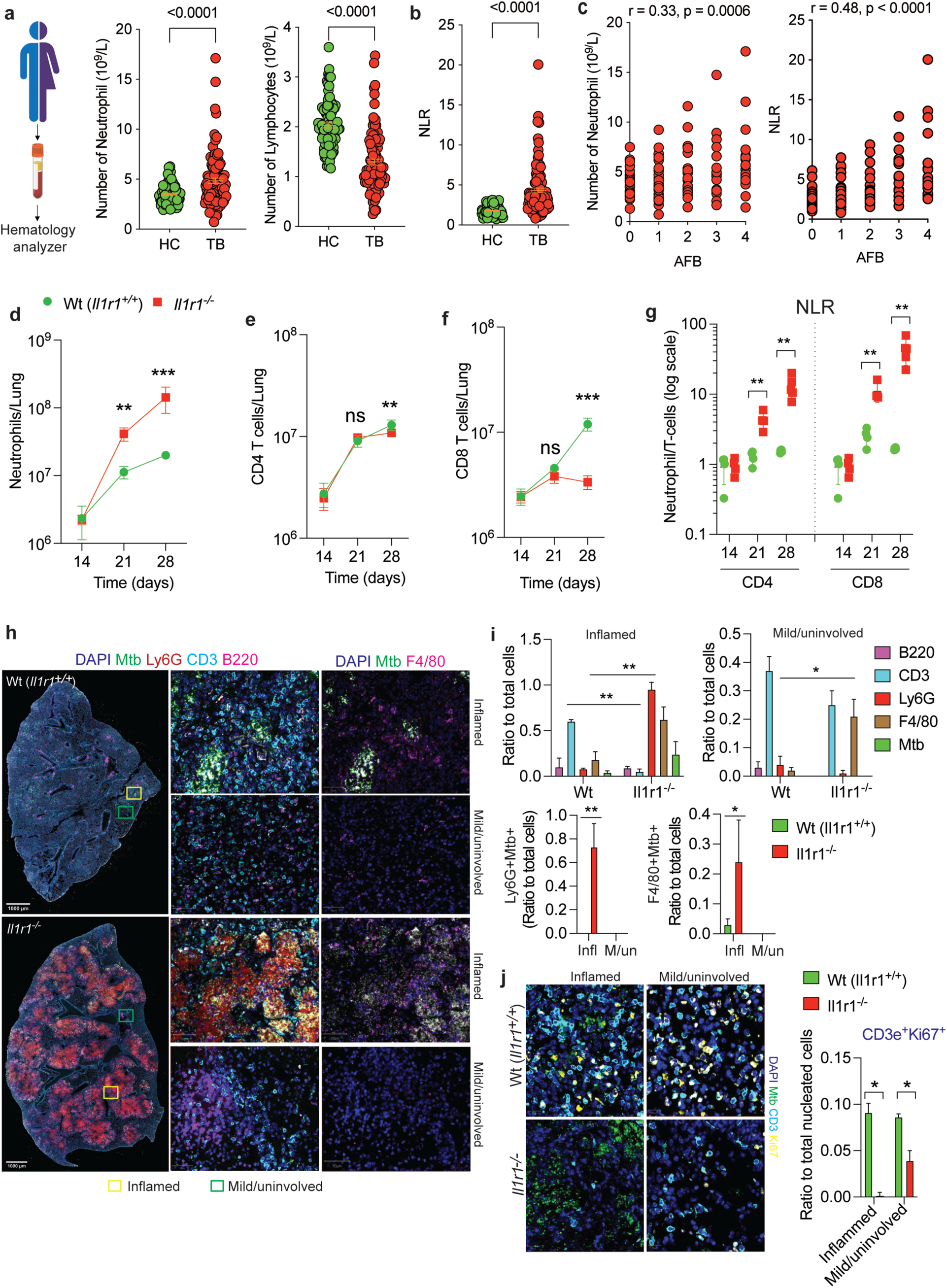
Abundance of infected neutrophils in TB lesions impairs T-cell immunity. (**a**) Number of neutrophils and lymphocytes in the peripheral blood of healthy (HC) and pulmonary TB patients (TB); (**b**) Ratio of absolute number of neutrophils to lymphocytes (NLR) in the peripheral blood of HC and TB. (**c**) Correlation of the number of neutrophils (left) or NLR (right) with AFB score in TB. AFB were detected by Ziehl-Neelsen staining; (a-c), HC, N =100, TB, N=102. Data are expressed as Mean ± SEM, an unpaired *t*-test was used to analyze differences between two groups. P-values was shown in the top of each panel. (**d-f**) Wild type (*Il1r1^+/+^*) and *Il1r1^−/−^* mice were infected with Mtb HN878, sacrificed at indicated time points post infection, infiltration of neutrophils and T (CD4, CD8) cells were determined by flow cytometry. (**g**) Neutrophils to CD4 and CD8 T-cells ratio at indicated dpi. Sample size of N=4-6 mice were included for each strain for each time point. Data are presented as Mean±SD, *, p<0.05; **; p<0.01, Two-way ANOVA with Bonferroni post-test. Representative of two individual experiments. (**h**) Cell DIVE multiplexed immunofluorescence (MxIF) was used to determine the abundance of neutrophils (Ly6G), T (CD3) and B (B220) cells, macrophages (F4/80) and bacteria (Mtb) in the lung of indicated strains of mice. Representative MxIF images from the FFPE lung sections, and region of interests randomly selected from inflamed and mildly inflamed/uninvolved regions of the lung from indicated strains of mice show the infiltration of immune cells and presence of bacteria. (**i**) Quantitation of the immune cells, infected myeloid cells in region of interest (ROI) (N=3 each) across mouse genotypes. (**j**) Representative MxIF images and quantitation of proliferating CD3e^+^T-cells in the ROIs of wt and *Il1r1^−/−^* mice lung. Data are presented as Mean±SD, statistical significance was calculated by One-way ANOVA with Sidak’s multiple comparison test, *, p<0.05; **; p<0.01. Source data are provided as a Source data file.

We next determined the kinetics of neutrophil and T-cell infiltration in the lung of wild type BL/6 (wt) and *Il1r1^−/−^* mice after infecting them with a reporter strain of Mtb HN878 expressing YFP. Neutrophils persistently increased in the *Il1r1^−/−^* mice compared to wt where a modest rise was observed. In contrast, both the CD4 and CD8 T-cells were elevated in wt mice as infection progressed. Strikingly, with increasing number of neutrophils, a proportionate decline in CD4 and CD8 T-cells were detected in the lung of *Il1r1^−/−^* mice. The distinct infiltration pattern of neutrophils and T-cells in both these strains was manifested as a relatively constant NLR for wt mice, while a rising NLR in the susceptible *Il1r1^−/−^* mice with disease progression (**Fig. 1d-g**). The escalation in NLR was associated with failed bacterial control in the lung as evidenced by higher bacterial load in myeloid cells, predominantly neutrophils (**Extended data Fig. 1i**).

To obtain insights into spatiotemporal organization of neutrophils and lymphocytes (CD3e^+^T cells and B220^+^ B-cells) in TB lesions, we performed multiplexed immunofluorescence (MxIF) staining of the infected lungs by employing Cell DIVE™, an reiterative staining and imaging workflow that allows spatial analysis of different cell types, including immune cells at a single cell resolution (**Extended Data Fig. 1j**)^32,33^. Consistent with previously published studies in mice and NHPs^34,35^, we found iBALT-like structures containing B220^+^ and CD3e^+^ T-lymphocytes mainly in the wt-mouse lung with fewer Ly6G^+^ neutrophils and F4/80^+^ macrophages. In contrast, Ly6G^+^ myeloid clusters were seen throughout the lung of *Il1r1^−/−^* mice. Importantly, these consolidated inflammatory structures had higher bacterial load (Mtb^+^), coinhabited by necrotic F4/80^+^ macrophages but excluded CD3e^+^ or B-220^+^ lymphocytes (**Fig. 1h**). We measured the ratio of the fluorescence counts of each biomarker relative to the total counts in a defined region of interest (ROI). CD3e^+^ cell counts in both the “inflamed” and “mildly inflamed/uninvolved” regions of the lung of *Il1r1^−/−^* mice markedly reduced compared to wt. Strikingly, neutrophil-rich inflammatory consolidations were the major infected sites in the lung with higher proportion of Ly6G^+^Mtb^+^ and F4/80^+^Mtb^+^ cells in *Il1r1^−/−^* mice compared to wt lung. Of note, the B220, F4/80 expressing cells were comparable between the wt and *Il1r1^−/−^* mice in the uninvolved regions, but fewer CD3e^+^ T-cells were detected in the latter (**Fig. 1i**). Next, we determined the proportion of proliferating CD3e^+^ T-cells by quantifying Ki67 expression, a prototypic biomarker of cell proliferation. No effect of inflammation on T-cell proliferation was observed in the wt mice. In stark contrast, Ki67^+^ CD3e^+^ cells were hardly detected within and around the neutrophil-rich inflamed regions of *Il1r1^−/−^* mice. Although these cells were detected in the regions with mild or no inflammation, they were significantly less compared to corresponding regions in wt mice (**Fig.1j**). Collectively, the kinetics of disease progression, and comprehensive spatial analysis of immune cells in the infected lungs provided a compelling indication that neutrophil dominance in TB lesions negatively influences T-cell response, causes lung damage, bacterial growth and consequently increases susceptibility to Mtb infection.

### Neutrophil depletion restores T-cell immunity

We and others have previously reported that neutrophils can serve as permissive niche for Mtb. As neutrophils were the predominant myeloid cells harboring bacteria in the lung of *Il1r1^−/−^* mice (**Extended Data Fig.1i**), we depleted neutrophils by administering anti-Ly6G (clone 1A8) antibody to *Il1r1^−/−^* mice (**Extended Data Fig. 2a, b**) and evaluated its impact on T-cell response and disease outcome. Neutrophil depletion ameliorated pulmonary damage, prevented weight loss, reduced bacterial load by >2.5 log_10_, and restored CD4 and CD8 T-cell numbers to wt level in both lung and spleen (**Fig. 2a-e**). The percentage of T-cells expressing Ki67 was significantly higher in *Il1r1^−/−^* mice lung as compared to wt **(Fig. 2d)**. Additionally, lung T cells across the genotypes, produced similar level of IL-2, a cytokine required for T cell proliferation **(Extended Data Fig. 2e)**. In contrast, the percentage and number of T cells expressing Ki67 or IL-2 secretion by T cells in the spleen of *Il1r1^−/−^* mice was markedly lower than the wt. Strikingly, T-cell regained their ability to proliferate and secrete IL-2 following 1A8 treatment **(Fig. 2d-e, Extended Data Fig. 2c-e)**. Based on these observations, we concluded that impaired T-cell functions is a consequence of systemic effects of neutrophils instead of direct MDSC like local immunosuppression in the lung.

**Fig. 2.**
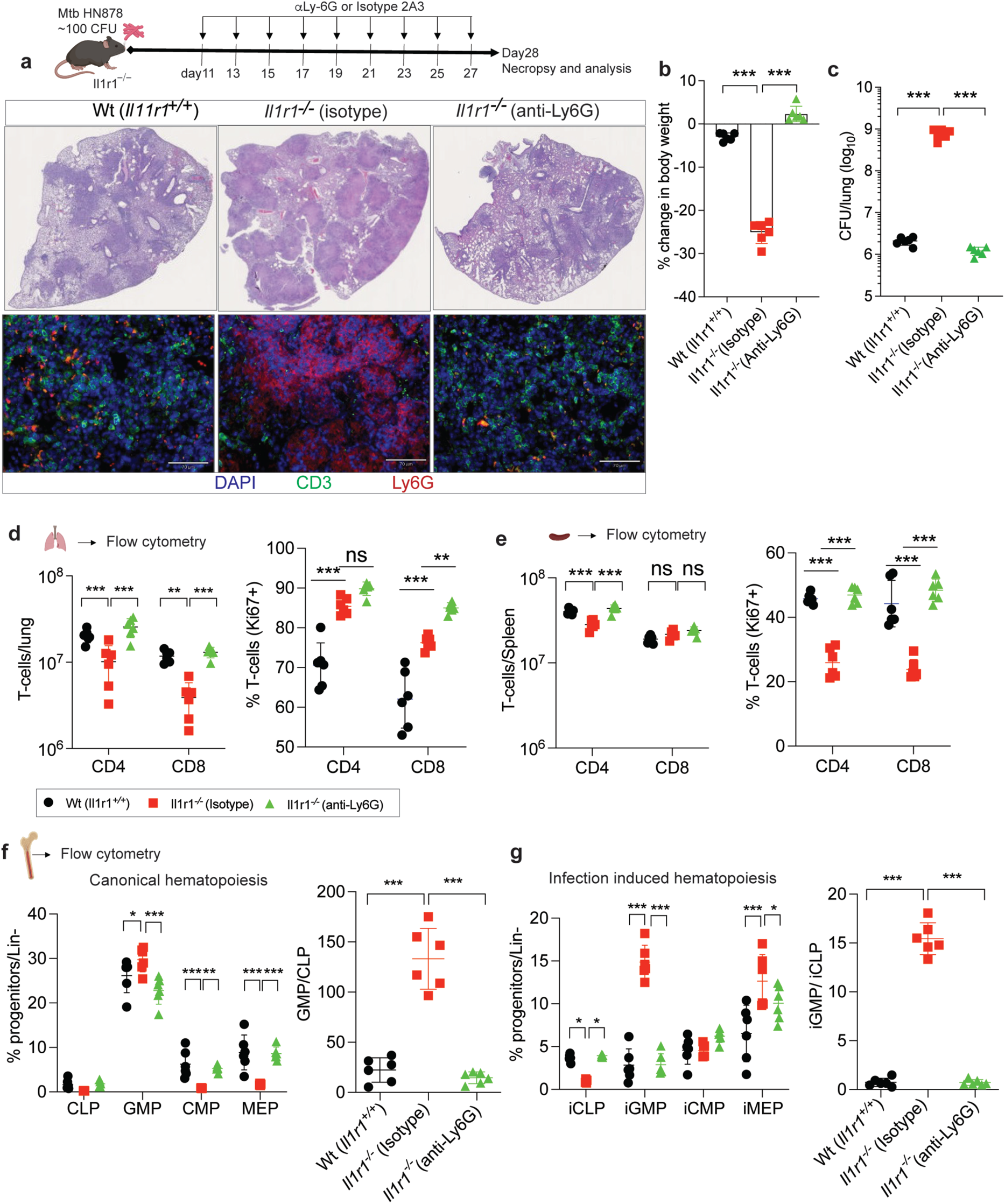
Neutrophil depletion restores T-cell immunity. Wild type (*Il1r1^+/+^*) and *Il1r1^−/−^* mice were infected with ~100 CFU of Mtb HN878. *Il1r1^−/−^* mice were treated with anti-Ly6G or isotype (N=5 mice each) shown in the schematics. At 28 dpi, mice were sacrificed. Untreated wild type (*Il1r1^+/+^*) mice were included in all experiments to determine the base line metrics of disease (**a**) Representative images of FFPE lung sections showing hematoxylin and eosin-staining (top panel) and immunofluorescence staining for neutrophils (Ly6G) and T-cells (CD3) in indicated cohorts (bottom panel). (**b**) Percentage of weight loss and (**c**) Bacterial load determined by CFU assay. **, p<0.01; ***, p<0.00, one-way ANOVA with Bonferroni multiple comparison test. (**d**) At 28 dpi, total number of CD4- and CD8-T-cells, percentage of proliferating T-cells (expressing nuclear antigen Ki-67) in the lung and (**e**) spleen were determined by flow cytometry. (**f**) Frequency of hematopoietic progenitors, relative abundance of GMP over CLP depicting canonical and (**g**) infection-induced hematopoietic response are shown in the linage^−ve^ cells of the bone marrow of indicated cohorts. CLP: common lymphoid progenitors, GMP: granulocyte macrophage progenitor, CMP: common myeloid progenitor, and MEP: megakaryocyte erythrocyte progenitor. Data are presented as Mean±SD. Significance was calculated by two-way ANOVA with Bonferroni multiple compassion test (d-g) and one way-ANOVA with Bonferroni multiple comparison test (f-g). *, p<0.05; **; p<0.01; ***; p<0.001. All the data are representative of two independent experiments.

To determine if neutrophil depletion has any impact on hematopoiesis that potentially could alter T-cell production in the bone marrow (bm) and ultimately in the peripheral tissue, we quantified the lineage committed progenitors in the bm. During hematopoiesis, granulocyte macrophage progenitors (GMPs) and common myeloid progenitors (CMPs) give rise to neutrophils and major myeloid cells. Common lymphoid progenitors (CLPs) are committed to produce T and B lymphocytes and megakaryocyte erythroid progenitors (MEP) produce platelets and red blood cells^36^. Mtb infection induced the expression of the stem cell antigen Ly6A/Sca1 on lineage^−ve^ cells in the bm. Thus, we categorized the Lin^−^cKit^+^ progenitors into Sca1^−/low^ and Sca1^hi^ populations. The Lin^−^cKit^+^Sca1^−/low^ and Lin^−^cKit^+^Sca1^hi^ cells were used to quantify various progenitors induced by “canonical hematopoiesis” and “infection-induced or demand-adapted hematopoiesis” (**Extended Data Fig. 3**). Mtb HN878 infection increased the frequency of GMPs, and infection induced GMPs (iGMP) among the lin^−^ bm cells in *Il1r1^−/−^* mice compared to *wt* (**Fig 2f, g**). The higher frequency of GMPs and iGMPs was associated with a decline in CLPs and infection induced CLPs (iCLPs). However, 1A8-treatment in *Il1r1^−/−^* mice resulted in significant reduction of GMP and iGMP frequency, whereas the CLPs and iCLPs frequency were restored to *wt* level (**Fig 2f, g**). Next, we quantified the relative abundance of GMP verses CLP and iGMP verses iCLP in the total live bm cells and found an elevated GMP/CLP and iGMP/iCLP ratio in the susceptible *Il1r1^−/−^* mice that was restored to wt equivalent upon neutrophil depletion (**Fig. 2f, g**). The high GMP/CLP ratio represents a “granulocyte-biased hematopoiesis” and iGMP/iCLP ratio depicts an “Mtb-infection-induced emergency granulopoiesis” response during TB in mice. To determine if this ratio could predict the susceptibility in mice from Mtb exposure, we infected wt BL/6, *Il1r1^−/−^*, *iNos^−/−^*, and C3HeB mice with Mtb HN878 and measured this ratio. Subsequently, we compared the GMP/CLP and iGMP/iCLP ratio in the bm of mice infected with Mtb HN878 to animals infected with relatively less virulent strain H37Rv and strains lacking type VII secretion system (Δesx1), to test the relevance of this readout to bacterial virulence. Mtb strain lacking esx1 (Δesx1) was less effective in inducing a GMP-biased hematopoiesis and preferentially induced more CLP production. Of note, no iGMP and iCLP was detected owing to its inability to induce Ly6A/Sca1 expression in lin^−^ bm cells (**Extended Data Fig. 4a, b**). The GMP/CLP ratio was higher in the bm of Mtb HN878 infected *Il1r1^−/−^*, *iNos^−/−^*, and C3HeB mice compared to BL/6 and depended on the virulence of the Mtb strain (**Extended Data Fig. 4c, d**). Together, these data indicated that neutrophil infiltration in the lung negatively impacted T-cell response by promoting hematopoietic remodeling in the bone marrow that preferentially produces more neutrophils at the expense of T-lymphocytes.

### Neutrophil derived G-CSF negatively regulates T-cell response

Cytokine analysis of Mtb infected lung lysates revealed the elevated level of cytokines (IL-1α, β), chemokines (MCP-1, MIP-1α, MIP-1β) and granulocyte colony stimulating factor (CSF3/G-CSF) implicated in neutrophil recruitment, growth, and survival respectively in the *Il1r1^−/−^* mouse lung during disease compared to wt and were significantly reduced upon neutrophil depletion. Interestingly, G-CSF concentration in the lung was almost undetectable in the neutrophil depleted *Il1r1^−/−^* mouse lung compared to isotype treated animals (**Fig. 3a**). Because neutrophil depletion affected hematopoiesis in the bone marrow (**Fig 2f, g**), we postulated that G-CSF, being a granulopoietic cytokine probably governs this phenotype. To examine if neutrophils serve as the source of this cytokine during Mtb infection, we analyzed lung sections by immunofluorescence staining. Although neutrophils from both the genotypes produced G-CSF, the higher abundance of these cells in the *Il1r1^−/−^* lung may account for the ~100-fold more G-CSF detected in the latter. Remarkably, 1A8 treatment almost abrogated the expression of G-CSF in the *Il1r1^−/−^* lung consistent with the cytokine quantitation in the organ homogenates (**Fig. 3b**). In congruence with the observation in mice, higher concentration of G-CSF was also detected in the plasma of individuals with active pulmonary TB (ATB) compared to latent tuberculosis infection (LTBI) and healthy cohorts (**Fig. 3c**).

**Fig. 3.**
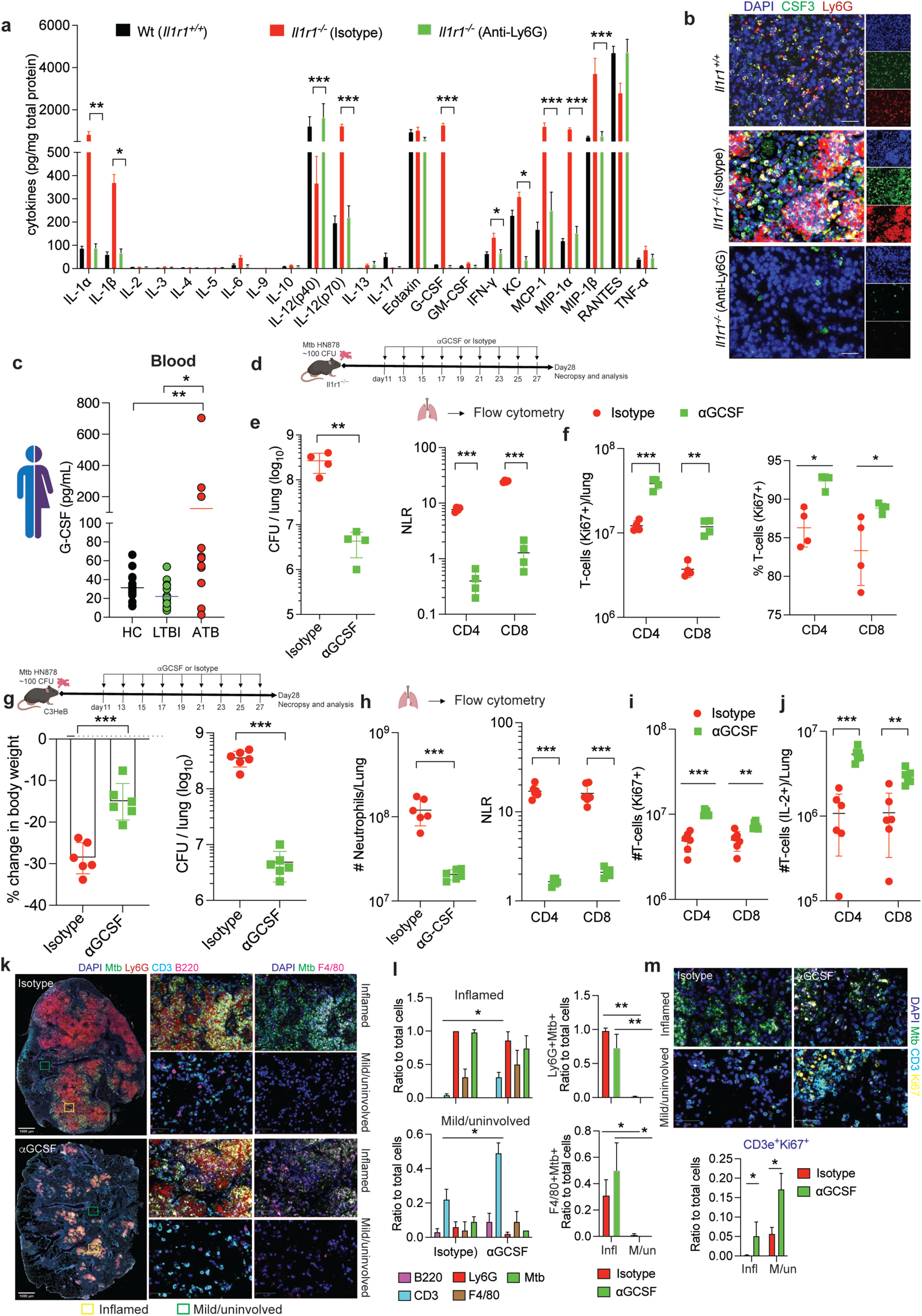
Neutrophil derived G-CSF negatively regulates T-cell response. (**a**) Cytokines in the lung homogenates of indicated mice were measured and normalized to the total protein content of each sample. (**b**) Representative immunofluorescent images of lung showing neutrophils (Ly6G) expressing G-CSF/CSF3. (**c**) The concentration of G-CSF was measured in the plasma of patients with HC (20), TB (n=20) and LTBI (n=17). Significance was calculated by (a) two-way ANOVA with Bonferroni multiple comparison test and (c) unpaired two-tail-student’s *t* test. *p < 0.05, **p < 0.01. (**d**) *Il1r1^−/−^* mice were infected with Mtb HN878, treated with anti-mouse G-CSF or isotype (N=5 mice each) as shown in the schematics. At 28 dpi, mice were sacrificed, (**e**) bacterial burden by CFU (left) and, NLR (right), and (**f**) number (left) and percentage (right) of proliferating T-cells (Ki67^+^) in the lung were determined. Data are presented as Mean±SD. Significance was calculated by two-way ANOVA with Bonferroni multiple comparison test. *, p<0.05; **; p<0.01; ***; p<0.001. The data are representative of two individual experiments. (**g**) C3HeB/FeJ mice were infected with Mtb HN878 and treated with anti-G-CSF (αG-CSF) or isotype as shown in the schematics. At 28 dpi mice were sacrificed, % weight loss (left) and bacterial burden (right) in the lungs were measured in these cohorts. (**h**) neutrophil number (left) and NLR (right), number of (**i**) proliferating and (**j**) IL-2 secreting T-cell subsets was shown. Sample size of N=6 was kept for each treatment group. Statistical significance was measured as shown in (e) and (f-g) respectively. (**k**) Representative Cell DIVE MxIF mages from the whole lung, and region of interests randomly selected from inflamed and mildly inflamed/uninvolved regions of the lung from both treatment groups of mice show the infiltration of immune cells and presence of Mtb. (**l**) Quantitation of the immune cells and Mtb infected myeloid cells in inflamed and mildly inflamed/uninvolved ROIs (n=3 each) across treatment groups. (**m**) Representative images (top) and quantitation of proliferating CD3e^+^ T-cells (bottom). Data are presented as Mean±SD, Statistical significance was calculated by One-way ANOVA with Bonferroni multiple comparison test, *, p<0.05; **; p<0.01. Source data are provided as a Source data file.

To determine whether G-CSF alters the protective T-cell response by promoting neutrophilic inflammation during TB, we analyzed the immune response in *Il1r1^−/−^*mice following G-CSF blockade by an αG-CSF neutralization antibody (**Fig. 3d**). G-CSF blockade led to >2 log_10_ and >1 log_10_ reduction in bacterial colony forming units (CFU) and neutrophil counts respectively compared to the IgG_2_ isotype treated cohorts (**Fig. 3e, Extended Data Fig. 5a**). The effect of G-CSF neutralization was particularly apparent on neutrophils and T-cells as evidenced by a significant reduction in NLR, while other myeloid and lung cells (CD45^+^CD11b^+^) were comparable between αG-CSF and isotype-treated *Il1r1^−/−^* mice (**Fig. 3e, Extended Data Fig. 5a**). G-CSF blockade significantly reduced the neutrophil influx (**Extended Data Fig. 5a, b**) and concomitantly enhanced the proliferating (Ki67^+^) CD4 and CD8 T cells in both lung and spleen (**Fig. 3f, g; Extended Data Fig. 5c**).

G-CSF is known to promote Th2-biased immune response in murine models of cancer^37,38^. Since Th1 immunity is critical for host defense against mycobacteria^26^ and Mtb HN878 impairs this function^39^, we asked if G-CSF negatively sculpted T-cell differentiation, and function. We measured the expression of lineage-specific transcription factors and cytokine secretion in CD4 and CD8 T-cells by intracellular staining for respective antigens. We observed a significant decline in T-bet, and Rorγt expression and a corresponding effect on IL-2, IFN-γ and IL-17 secretion by CD4, CD8 T cells in the spleen of *Il1r1^−/−^* mice compared to wt animals. Remarkably, αG-CSF treatment restored their expression and augmented cytokine secretion from CD4 and CD8 T-cells either equivalent or higher than wt mice (**Extended Data Fig. 5d-e**). Notably, the effect of G-CSF on cytokine secretion was more evident in the spleen compared to lung, although αG-CSF treatment enhanced the cytokine secretion from CD4 and CD8 T-cells at a single cell level in the lung (**Extended Data Fig. 5e**). Mtb infection did not impact the expression of Gata3 in the splenic CD4 and CD8 T cells, and G-CSF inhibition led to a modest increase. However, a corresponding change in the prototypical Th2 cytokines, IL-4, and IL-10, was not observed in the lung and spleen before and after G-CSF blockade (data not shown). These data suggest that overproduction of G-CSF has profound negative effects on the systemic Th1 and Th17 responses during Mtb infection that are critical for anti-mycobacterial immunity.

To determine if the G-CSF mediated alteration of T-cell immunity is a general mechanism of pathogenesis during TB, we repeated the G-CSF neutralization in C3HeB mice (**Fig. 3g**). G-CSF blockade significantly ameliorated disease as manifested in significant reduction of weight loss, bacterial burden (~2 log_10_), neutrophil influx to the lung (**Fig. 3g, h**). Coherent with the observations in *Il1r1^−/−^*mice, αG-CSF treatment resulted in significant increase in the Ki67^+^ and IL-2^+^ T-cells in the lung and spleen (**Fig. 3i-j**, **Extended Data Fig. 5f**). We leveraged Cell DIVE MxIF to gain further insights into the spatial relationship of neutrophils and T-cells in the Mtb infected lung with and without G-CSF inhibition. As observed in *Il1r1^−/−^* mice, inflammatory consolidations containing infected neutrophils (Ly6G^+^Mtb^+^) was a pathological hallmark of Mtb HN878 infected lung in C3HeB mice. Very few CD3e^+^ T-cells were detected within the myeloid cell core containing the bacilli. However, G-CSF blockade significantly reduced the number of neutrophil-rich myeloid clusters in the lung, and increased total (CD3e^+^), and proliferating T-cells (Ki67^+^) compared to isotype treated mice. The effect of G-CSF inhibition was less pronounced on B220^+^ and F4/80^+^ cells. Notably, G-CSF inhibition also increased the overall T-cell numbers and proliferation in the regions of lung that were selected based on mild or no inflammatory involvement (**Fig. 3k-n**). G-CSF mediated effects on T-cells was not dependent on receptor engagement on these cells as inhibition of G-CSFR by anti-CSF3R antibody did not affect proliferation (Ki67^+^) or cytokine secretion (IL-2, IFN-γ secretion) of CD4 and CD8 T-cells (**Extended Data Fig. 5g-j**).

Next, we determined if G-CSF acts on neutrophils rendering them immunosuppressive functions that underlies its inhibitory effects on T-cell response. By using a T-cell, neutrophil co-culture system described before^40^, we quantified T-cell proliferation as our final readout with and without recombinant murine G-CSF (rG-CSF) stimulation. G-CSF stimulation of neutrophils was unable to inhibit T-cell proliferation (**Extended Data Fig. 6a**). Taken together, these observations implied that neutrophil infiltration during Mtb infection negatively influenced T-cell immunity by a G-CSF dependent mechanism that do not involve MDSC-like functions of these cells, and G-CSF may not directly mediate these effects on T-cells.

### G-CSF promotes a granulocyte-biased hematopoiesis to suppress T-cell response

G-CSF is the principal hematopoietic cytokine regulating granulopoiesis and is known to direct the expansion and differentiation of hematopoietic stem cells (HSCs) to skew hematopoiesis towards the myeloid lineage. Chronic skewing of myeloid production could affect the abundance of other immune cells including T-cells by impacting common lymphoid progenitors (CLPs). Based on these reports, we hypothesize that G-CSF regulates T-cell response by modulating hematopoiesis in bm. To test this, we administered rG-CSF to naiive C3HeB mice and measured various hematopoietic progenitors (**Extended Data Fig. 6b, c**). Consistent with the previous reports^41^, rG-CSF administration specifically increased the frequency and number of GMPs and CMPs with a concomitant decline in the CLPs (**Extended Data Fig. 6c**). The increase in GMPs was reflected in the higher numbers of neutrophils in the bm, blood, spleen, and lung compared to vehicle treated mice (**Extended Data Fig. 6d**). In contrast, rG-CSF administration caused a decrease in the percentage of proliferating CD4, CD8 T-cells in both the lung and spleen (**Extended Data Fig. 6e, f**). It is notable that rG-CSF administration did not alter Ly6A/Sca1 expression in lin^−^ cells that was induced during Mtb infection (**Fig. 2f, g**). Given the effect of rG-CSF in driving granulopoiesis, we next asked if G-CSF blockade during Mtb infection inhibits emergency granulopoiesis, and enriches CLPs in the bm. Although G-CSF blockade did not significantly alter the frequency of GMPs or iGMPs and CLPs or iCLPs (**Extended Data Fig. 6g**), it markedly reduced the immature neutrophils (myelocytes, metamyelocytes) and augmented the mature cells (**Extended Data Fig. 6h, i**). Together, these results indicate that G-CSF plays a crucial role during steady state hematopoiesis resulting in the accumulation of immature cells in the bm. However, during infection, G-CSF probably collaborates with other pathways to regulate infection-induced hematopoiesis that disrupts the balance of GMP and CLP production.

### Type I IFN pathway genes are enriched in the neutrophils from different organs

Neutrophils are generated in the bm, mobilized to the circulation, and then recruited to the lung in response to infection. Since neutrophils serve as a conducive niche for the bacteria^21,40^, it is likely that Mtb exploits this innate response by inducing the synthesis, mobilization, and survival of neutrophils in infected organs. We therefore sought to identify pathways/factors that are specifically induced in neutrophils derived from *Il1r1^−/−^* mice that produced pathologic level of G-CSF compared to their wt counterparts. We purified neutrophils from the lung, spleen, and bm of wt and *Il1r1^−/−^* mice and analyzed their transcriptome by bulk mRNA sequencing. Principal component analysis of the differentially expressed genes (DEGs) showed that lung, spleen, and bm neutrophils of these mice were separately clustered (**Fig. 4a**). Differential gene expression analysis in these organs uncovered that, 987 genes were upregulated, and 1210 genes were downregulated in the lung of *Il1r1^−/−^* mice compared to wt neutrophils. Similarly, in the spleen and bm, 781 and 507 genes respectively were upregulated, and 1052 and 1021 genes were downregulated in *Il1r1^−/−^* mice compared to wt neutrophils (**Extended Data Fig. 7a, Supplementary Table 2**).

**Fig. 4.**
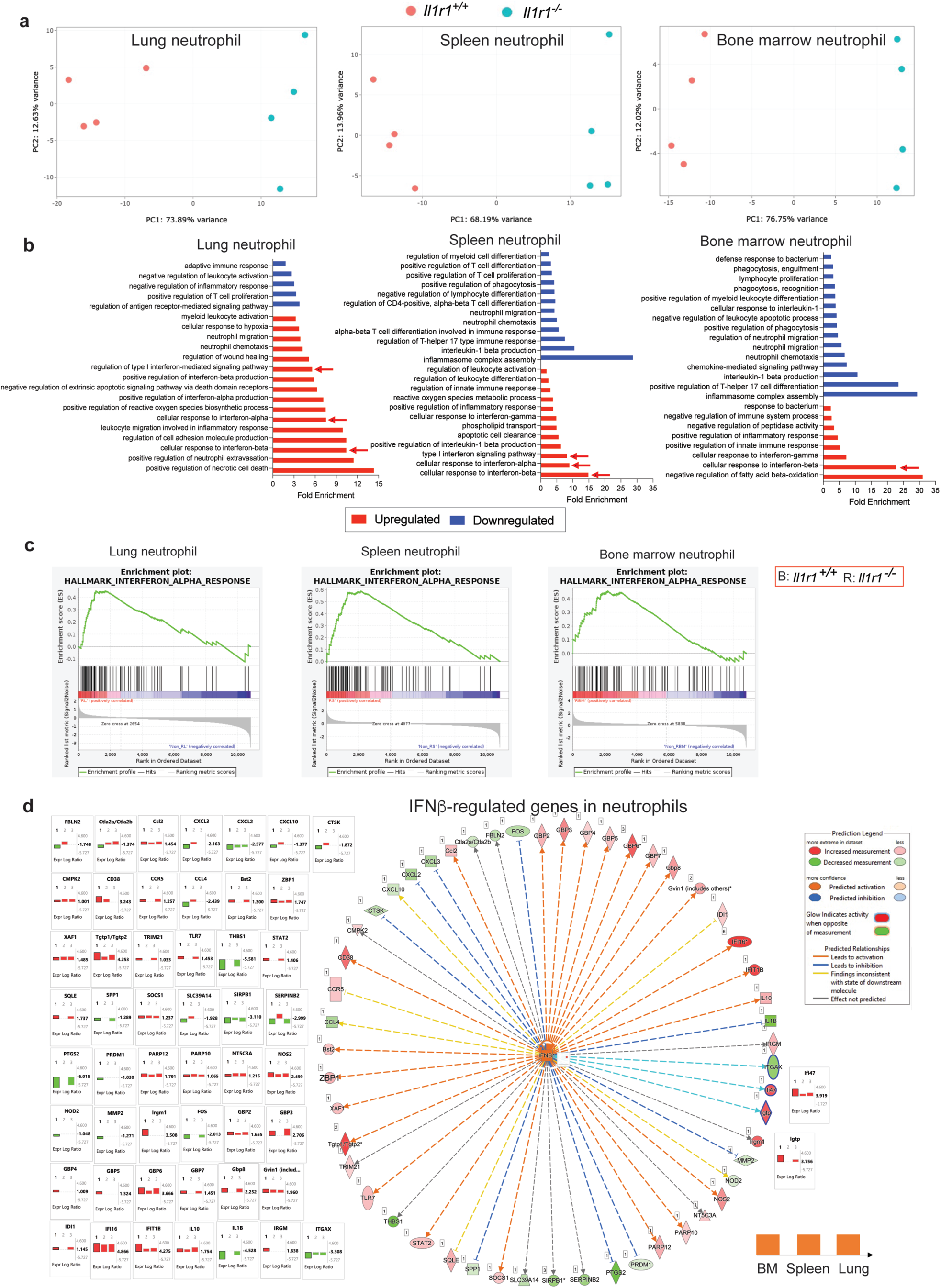
RNA sequencing of neutrophils reveals upregulation of type I IFN regulated genes and pathways to associate with TB pathogenesis. Wild type (*Il1r1^+/+^*) and *Il1r1^−/−^* mice were infected with Mtb HN878. At day 25 dpi, neutrophils purified from lung, spleen and bone marrow single cell suspensions were processed for mRNA sequencing (see methods). (**a**) Co-variance based principal component analysis scatter plots shows separate clustering for different groups (N=4 each). (**b**) Bar graphs demonstrate the up-regulated and downregulated genes belonging to pathways in neutrophils of *Il1r1^−/−^* mice in each tissue compared to the wild-type (*Il1r1^+/+^*) neutrophils. Arrows indicate the upregulation of gene sets belonging to the type I IFN (interferon-alpha and beta) pathways. (**c**) GSEA analysis shows the enrichment of genes implicated in interferon-α response in neutrophils across organs. (**d**) Ingenuity software based custom network view of all genes directly (solid lines) and indirectly (dashed lines) associated with interferon-β in neutrophils from *Il1r1^−/−^* mice versus wild-type (*Il1r1^+/+^*) across organs. Upregulated genes are shown in shades of red and downregulated genes in shades of green.

WGCNA analysis of all the genes across the three organs identified gene modules previously reported in Mtb HN878 infected mouse lung^42^ and peripheral blood of human TB patients^24^. Among the 15 common gene modules previously reported in human TB^18^, the “interferon/PRR module” was highly enriched in the neutrophils of both wt and *Il1r1^−/−^*mice, with high enrichment score recorded for *Il1r1^−/−^* mouse lung neutrophils (**Extended Data Fig. 7b**). We next performed a neutrophil pathway enrichment analysis in the RNAseq DEG dataset obtained from different organs. 145 neutrophil specific pathways were selected from the molecular signatures database (MSigDB C2 curated pathways, see Methods) to determine how many pathways in our dataset were enriched in the neutrophils from three organs to identify common gene modules. Raw GSVA enrichment score was used to generate dot plots for the 22 pathway modules, displaying significant (p<0.05) differences among *Il1r1^−/−^* mice and wt. This analysis again identified enrichment of IFN-I modules in the neutrophils from the bm, spleen and lung with higher enrichment score for lung neutrophils (**Extended Data Fig. 7c, indicated boxes**).

Gene ontology (GO) analysis of the DEGs in the neutrophils revealed that these cells exhibit tissue specific signatures. However, among the common GO pathways overrepresented in the neutrophils of *Il1r1^−/−^* mice compared to wt, belonged to the “cellular response to interferon β” and “cellular response to interferon α/β” in the lung, spleen and bm (**Fig. 4b, Extended Data Fig. 8a**), suggesting that neutrophils from these organs despite exhibiting different transcriptional signatures, respond to IFN-I by expressing genes related to type I IFN signaling pathways. GSEA analysis of DEGs revealed that in addition to genes related to innate immune response, neutrophil chemotaxis, IFNγ response, IFNα pathway regulated genes were enriched in the neutrophils from all organs (**Fig. 4c and Extended Data Fig. 8b**). Next, we performed ingenuity pathway analysis (IPA) in the DEG dataset from *Il1r1^−/−^* and wt neutrophils to determine the representation of IFN-I gene signatures that showed different ISGs (Igtp, Gbp2, Gbp4, Ifi16, Ifi47, Irgm1) were upregulated in the neutrophils of *Il1r1^−/−^* mice over wt though their log_2_-fold changes vary dramatically depending on the tissue microenvironment (**Fig. 4d, Extended Data Fig. 8b-d**). Collectively, this comprehensive transcriptome analysis of neutrophils suggests, these cells respond to the robust production of IFNα/β elicited by Mtb HN878 infection.

### Type I IFN receptor expression is associated with TB disease in humans

Responsiveness to IFN-I requires the expression of IFNAR on a target cell which is a heteromeric cell surface receptor composed of two subunits, referred to as the low affinity subunit, IFNAR1 and the high affinity subunit, IFNAR2^43^. Since IFN-I response signature is a hallmark of TB disease in humans, we analyzed the IFNAR1 and IFNAR2 expression in different publicly available transcriptome datasets of whole blood from active TB and healthy controls (HC). Neutrophils expressed highest level of IFNAR1 and IFNAR2 among the different cell types examined (**Fig 5a**). The expression of these receptors significantly increased in TB, compared to HC, other diseases (OD), and LTBI subjects (**Fig. 5b-d**). Next, we evaluated if IFNAR1 and IFNAR2 expression preceded progression to active disease. Expression levels of both the receptor subunits were significantly elevated in TB progressors, compared to non-progressors (**Fig. 5e**). Based on these observations in humans, and in-depth transcriptome analysis of mouse neutrophils, we concluded that IFN-I signaling response in neutrophils is critical for disease pathogenesis and perhaps contribute to hematopoietic regulation during TB.

**Figure 5.**
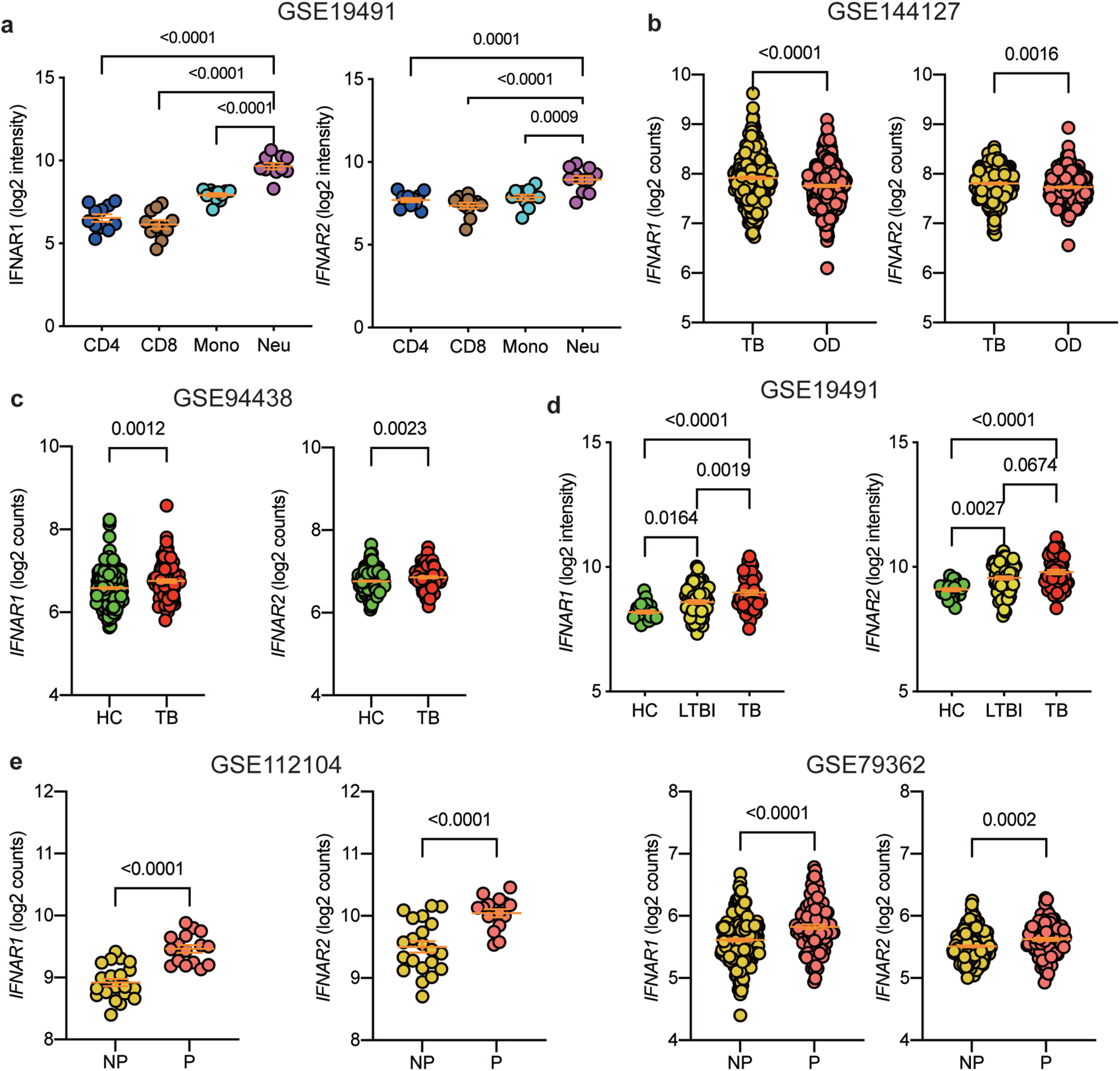
Increase expression of IFNAR1 and IFNAR2 are associated with human TB and disease progression. (**a**) IFNAR1 and IFNAR2 expression in different cell types. (**b**) IFNAR1 and IFNAR2 expression in active TB versus other disease (OD). (**c**) IFNAR1 and IFNAR2 expression in active TB versus HC, (**d**) LTBI and HC subjects. (**e**) IFNAR1 and IFNAR2 in non-progressor (NP) versus pro-progressor. A one-way ANOVA Newman–Keuls multiple comparison test was used to compare differences among multiple groups. An unpaired *t*-test was used to analyze differences between two groups. P-values was shown in the top of each panel.

### Type I IFN is essential for Mtb infection induced granulopoiesis

To examine the role of IFN-I in regulating the pathogenic granulopoiesis and impaired T-cell response following Mtb infection, we generated *Ifnar1* deficient mice in the *Il1r1^−/−^* background (*Il1r1^−/−^Ifnar1^−/−^*) and compared their susceptibility with the wt and *Il1r1^−/−^* mice. *Il1r1^−/−^Ifnar1^−/−^* mice were completely protected from Mtb HN878 challenge (**Fig. 6a**). The enhanced survival of *Il1r1^−/−^ Ifnar1^−/−^* mice coincides with significant reduction in both neutrophils and NLR in the lung (**Fig. 6b-d).** We generated mixed bone marrow chimeras by reconstituting *Il1r1^−/−^* recipient mice with 1:1 mix of wt (CD45.1) and *Ifnar1^−/−^* (CD45.2) bm cells and evaluated their hematopoietic response. Among the various progenitors examined, GMPs and iGMPs, were proportionately higher in *Ifnar1^+/+^*(wt) compared to the *Ifnar1^−/−^*donor cells. Notably, the latter had markedly higher frequencies of CMPs and MEPs. Moreover, *Ifnar1^−/−^* iCLPs and iCMPs were trending towards an increase, though it was not significantly different from wt donor cells (**Fig. 6e-g**). Cell-intrinsic Ifnar1 signaling impacted neutrophils at multiple nodes, spanning their synthesis, mobilization, and recruitment as evident by higher percentage of neutrophils among wt in bm, spleen and lung compared to *Ifnar1^−/−^* donor cells (**Fig. 6h-j**). The calculated NLR in the lung was higher for the wt-cells when compared to *Ifnar1^−/−^* cells (**Fig 6k**).

**Figure 6.**
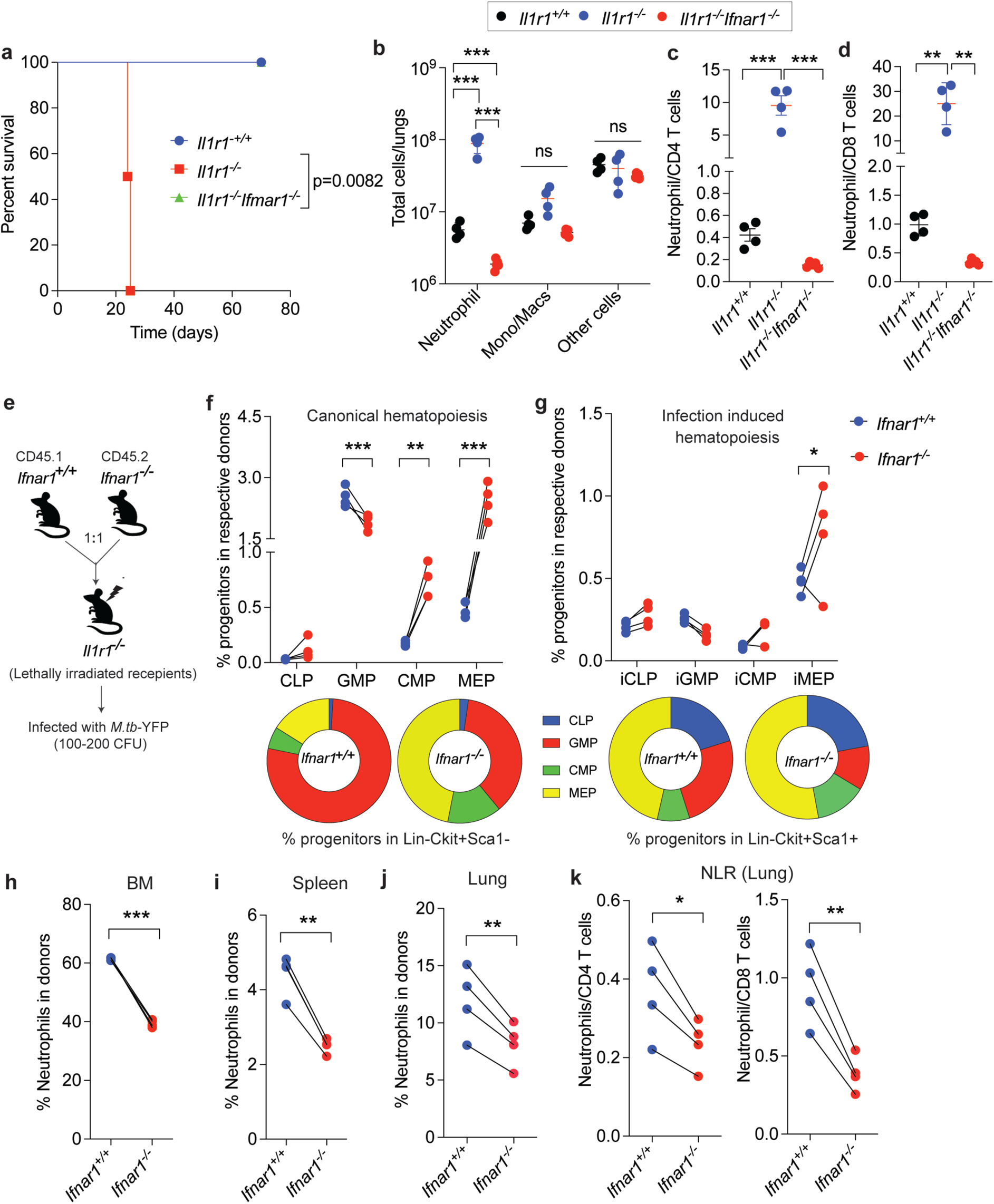
Type I IFN response is critical for promoting pathologic granulopoiesis and consequent neutrophilic inflammation. Wild type (*Il1r1^+/+^*), *Il1r1^−/−^* and *Il1r1^−/−^ Ifnar1^−/−^* mice were infected with Mtb HN878. (**a**) Overall survival was measured up to 70 dpi (N=8 mice/group) and shown as Kaplan-Meier plot. (**b**) Infiltration of immune cells, relative abundance of (**c**) neutrophils and CD4 T-cells and (**d**) CD8 cells in the lungs of these mice was shown. N=4 mice/genotype. Data are presented as Mean±SD, Statistical significance was calculated by (a) log-rank Mantel-Cox test, (b) two-way and (c, d) one-way ANOVA; Bonferroni multiple comparison test. *, p<0.05; **; p<0.01; ***, p<0.001. (**e**) Schematics of Mtb HN878 infection in mixed bone marrow chimeras. (**f, g**) Frequency of the lineage-committed progenitors expressing congenic markers within CD45.1 or CD45.2 bm cells (top) and Lin^−^ckit^+^sca1^−/low^ or Lin^−^ckit^+^sca1^+/hi^ (bottom donut) bm cells representing conventional, and infection induced hematopoiesis were shown. (**h**) Frequency of *Ifnar1^+/+^* and *Ifnar1^−/−^* neutrophils within the CD45.1 or CD45.2 cells in the bm, (**i**) spleen, (**j**) lung and (**k**) relative abundance of neutrophils and CD4 or CD8 T-cells in chimera mice was shown. Significance was calculated by two-way ANOVA with Bonferroni multiple comparison test. *, p<0.05; **; p<0.01. Source data are provided as a Source data file.

As Ifnar1 signaling is required for regulating pathological hematopoiesis and consequent neutrophil-dominant inflammation in *Il1r1^−/−^* mice, we then asked whether Ifnar1-regulated granulopoiesis is a general feature of exacerbating TB disease. We infected C3HeB mice with Mtb HN878-YFP, evaluated the hematopoietic response, and disease parameters after blocking Ifnar1 signaling by a neutralizing antibody reported before^16^. Compared to the isotype treated mice, Ifnar1 blockade significantly reduced the GMP/CLPs, iGMP/iCLPs, restored the frequency of CMPs and MEPs in the bm cells (**Fig. 7a-c**) as observed in *Il1r1^−/−^* mice. This alteration in hematopoiesis resulted in >3 log_10_ reduction in bacterial CFU in the lung, reversal of wasting disease, ~100-fold reduction in the number of infected neutrophils, and a concomitant increase in both CD4 and CD8 T-cells resulting in drastic reduction in NLR in the lung of αIfnar1 treated mice compared to control antibody treated animals (**Fig. 7d-i**). Single cell analysis by Cell DIVE MxIF demonstrated that Ifnar1 blockade reduced the overall neutrophil influx, myeloid cell foci containing infected neutrophils, and macrophages in the defined ROIs of lung (**Fig. 7j-k**). Most notably, IFN-I inhibition replenished the proliferating T cells (Ki67^+^CD3e^+^) in both inflamed and mildly inflamed/uninvolved ROIs (**Fig. 7l**). Consistent with the protective functions of T-cells, the higher abundance of CD3e^+^ T-cells over neutrophils was associated with a significant decline in the Mtb^+^ cell counts corroborating the bacterial burden measurements by CFU (**Fig. 7d**). Collectively, these data suggest that Ifnar1 signaling on neutrophils largely regulates the synthesis, mobilization, and recruitment of these cells to the lung and causes TB pathogenesis.

**Figure 7.**
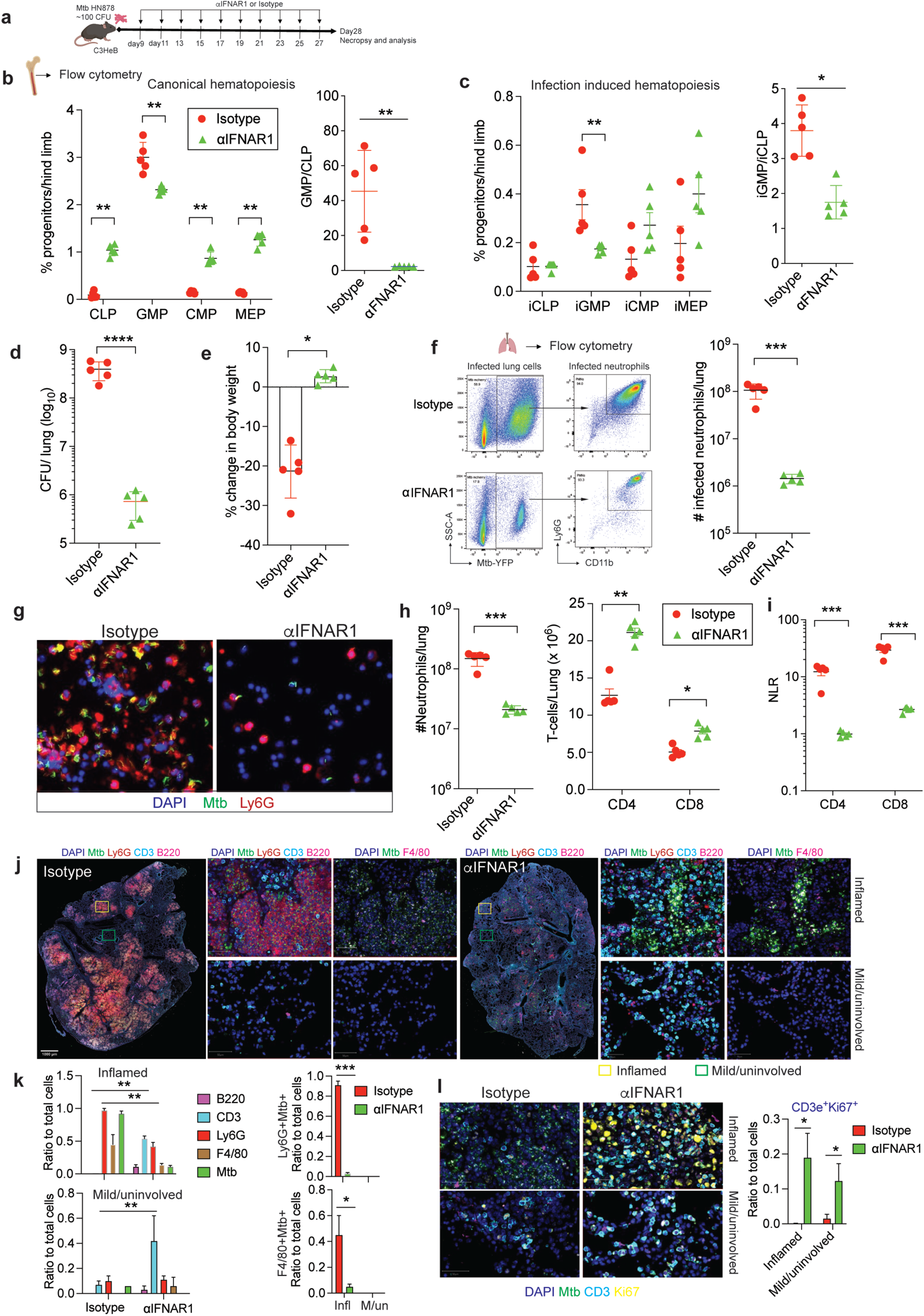
Inhibition of Ifnar1 signaling limits infection-driven pathogenic granulopoiesis. (**a**) C3HeB/FeJ mice were infected with Mtb HN878 (expressing YFP) and treated with anti-IFNAR1 or isotype (N=5 mice/group) as depicted in the schematics. At 28 dpi, mice were sacrificed, (**b**) frequency of lineage committed progenitors and relative abundance of GMP over CLP within the Lin^−^ckit^+^sca1^−/low^ and (**c**) Lin^−^ckit^+^Sca1^+/hi^ bm cells, depicting conventional and infection-induced hematopoiesis respectively were shown. Various metrics of disease was evaluated such as (**d**) bacterial load as CFU in the lung, (**e**) % change in body weight following treatment. (**f**) Representative FACS plots and quantification of total number of Mtb-YFP^+^ infected neutrophils in the lung and (**g**) representative immunofluorescent image of cytospins prepared from single cell suspension of lung showing Mtb-infected neutrophils (Ly6G+YFP+) in anti-IFNAR1 or isotype treated mice (N=5 mice/group). (**h**) Flow cytometry quantitation of neutrophils, CD4 and CD8-T-cells, and (**i**) relative abundance of neutrophil to CD4 or CD8 T-cells in the lung of anti-IFNAR1 or isotype treated mice. (**j**) Representative Cell DIVE MxIF images from the whole lung, and ROIs randomly selected from inflamed and mildly inflamed/uninvolved regions of the lung from both treatment groups of mice show the infiltration of immune cells and presence of bacteria. (**k**) Quantitation of the immune cells, infected cells in indicated ROIs (n=3 each) and (**l**) expression of the proliferation marker Ki-67 in CD3e^+^ T-cells across treatment groups. Data are presented as Mean±SD. A two-way ANOVA; Bonferroni multiple comparison test was used to compare differences among multiple groups and an unpaired *t*-test was used to analyze differences between two groups (b-i). *, p<0.05; **; p<0.01; ***, p<0.001, ****, p<0.0001. Source data are provided as a Source data file.

### Type I IFN is necessary and sufficient for pathologic granulopoiesis

Since Ifnar1 blockade ameliorated disease by limiting pathologic granulopoiesis, we determined whether IFN-I contributes to G-CSF-mediated inflammation, and consequent impairment in T-cell immunity during Mtb infection. To test this, we administered rG-CSF to mice that were previously treated with αIfnar1 or isotype control antibody (**Extended Data Fig. 9a).** As observed previously, αIfnar1 treatment significantly reduced the frequencies of GMPs, GMP/CLP and iGMP/iCLP ratio (**Extended data Fig. 9b, c**) compared to isotype treated animals. This alteration in hematopoiesis consequently led to a reduction in overall lung inflammation, total and infected neutrophils, increase in CD4 and CD8 T-cells in the lung, low NLR, and attenuation of histopathological disease compared to isotype treated animals (**Extended data Fig. 9e-i**). Notably, rG-CSF administration following αIfnar1 treatment did not exert an appreciable change to the hematopoiesis, neutrophil, and T-cell numbers or NLR in the lung (**Extended data Fig. 9e-i**). rG-CSF administration combined with Ifnar1 blockade profoundly increased the number of mature neutrophils in the bm (**Extended data Fig. 9d**), indicating that the G-CSF driven increase in mature neutrophils failed to mobilize from the bm in the absence of Ifnar1-regulated signals. Collectively, these results suggest that G-CSF may be required to sustain the neutrophil pool in the bm, but Ifnar1 signaling is both necessary and sufficient for mobilization and recruitment of neutrophils ultimately impairing T-cell responses during TB.

## Discussion

A hallmark of immune response to Mtb is the formation of organized multicellular structures called granulomas^1^. Cellular composition of granulomas in active TB disease and latent infection is relatively well characterized, but the molecular mechanism that underlies the formation of a protective or pathological lesion remains obscure. Our studies demonstrate that the neutrophils sculpt the granuloma fate by regulating T-cell positioning in TB lesions. Neutrophil depletion restores the number and spatial arrangement of T-cells in inflammatory lesions, resulting in bacterial containment and limiting immunopathology (Fig.2). In support of our mouse studies, high NLR in the peripheral blood was associated with pulmonary TB in humans and strikingly corelated with the severity of disease (Fig.1, Extended Data Fig 1). High NLR is known to differentiate active TB from bacterial pneumonia, sarcoidosis, associates with poor clinical outcomes^44^, predicts the risk of progressing to active disease in HIV-Mtb coinfection^45^ and mortality with development of acute respiratory distress (ARDS) in miliary TB^46^. Interestingly, high NLR is also reported in patients with severe COVID-19^47^ and poor prognosis from various cancers^48^. Our work highlights the role of G-CSF-IFN-I axis to regulate the balance between neutrophils and lymphocytes during chronic Mtb infection that can be targeted for rational designing of interventions for TB and possibly other respiratory infections.

Accumulating evidence suggests that neutrophils secrete several cytokines, chemokines, and growth factors during inflammation^49^ and in TB lesions of cynomolgus macaques^50^. In this work, we demonstrated that neutrophils function as bonafide producers of G-CSF, normally produced by endothelial cells, monocytes and macrophages, that not only fosters neutrophillic inflammation^51^ but inhibits CD4 and CD8 T-cell responses^38^. G-CSF is routinely used in clinical medicine to correct neutropenia associated with cancer chemotherapy^52^ and cytopenia following anti-TB antibiotics treatment^53^. The majority of allogeneic stem cell transplantation is currently performed by G-CSF-mobilized peripheral blood stem cells. Of note, G-CSF administration often associates with T-cell suppression in patients and requires direct G-CSFR signaling on these cells^54^. However, our results indicated that G-CSF mediated effects on T-cells during TB may not involve direct receptor engagement as G-CSFR antibody treatment did not alter proliferation or cytokine response in T-cells of the lung (Extended Data Fig.5). Instead, our data imply that G-CSF alters T-cell immunity during TB by promoting granulocyte expansion in the bone marrow that consequently suppresses lymphopoiesis. Since G-CSF possesses opposing effects on neutrophils and T-cells, persistent production of this cytokine during Mtb infection is potentially detrimental for the host. Indeed, G-CSF inhibition ameliorates neutrophil dominated pathology in *Card9^−/−^* mice and led to enhanced survival^55^. Our work in different susceptible mice supports these studies and provided mechanistic insights into the enhanced risk of developing disease from prolonged G-CSF production. Our work further argues against a generalized use of G-CSF for treating neutropenia as it may increase the risk of developing active disease in individuals with latent TB especially in TB endemic settings.

Airway concentration of G-CSF is positively corelated with neutrophillic inflammation during asthma^56^, lung metastasis^57^, Mtb infected inbred^40,58,59^, diverse outbred^60^ and genetically diverse collaborative cross mice^61^. In congruence with earlier studies, rG-CSF was able to mobilize neutrophils, and inhibit T-cell functions in naive C3HeB mice (Extended Data Fig 6). Conversely, G-CSF blockade by neutralizing antibody early during infection markedly reduced the neutrophil influx to the lung, and augmented T-cell functions in TB lesions (Fig. 3k). The abundance of Ly6G^+^ neutrophils in the inflamed regions of the lung with and without G-CSF inhibition was comparable, but the number of these lesions with myeloid cell consolidations were significantly reduced (**Fig.3k**). This reduction in the overall number of neutrophils, neutrophil clusters in the lung coincided with increased abundance of total and Ki67^+^ T cells in the inflamed areas. Importantly, the spatial arrangement of T-cells in the inflamed regions, and their enhanced recruitment to the lung resulted in fewer infected cells (Mtb+), further supporting studies in mice and NHPs that have previously shown T-cell positioning in TB lesions to be crucial for Mtb control through macrophage activation^6,27,62^. In comparison to G-CSF inhibition, neutrophil depletion had a more discernible impact on inflammation, bacterial control, suggesting that in addition to G-CSF production, neutrophils contribute to immunopathogenesis by an additional mechanism. mRNA sequencing analysis of neutrophils isolated from bone marrow, spleen and lung uncovered a role of type I IFN signaling response in neutrophils (Fig.4) to influence both the granulopoiesis and recruitment of these cells during TB.

Mtb infection induces rapid IFN-I production^63,64^ which is strain dependent^65^. Lineage 2 strains such as Mtb HN878 induces higher concentration of IFNα/β than the commonly used laboratory strains, and the virulence mechanism is linked to IFN-I-mediated impairment of Th1 immune response^39^. IFN-I has been shown to inhibit T-cell proliferation in vitro^66^ and delays Th1 response in the lung in a mouse model of LCMV-Mtb coinfection^67^. Here we showed that IFN-I inhibition in Mtb infected susceptible mice curtails neutrophil influx, augments T-cell proliferation, and most importantly improved the disease outcome. Additionally, our mixed bm chimera studies further indicated that cell-autonomous Ifnar1 signaling in neutrophils is critical for their trafficking, which coincides with alteration in T-cell responses in both the lung and spleen (Fig. 6). IFN-I has been shown to induce neutrophil-mediated pathology during TB by forming neutrophil extracellular traps (NETs) in the lungs^42^. Neutrophil intrinsic Ifnar1 signaling is therefore critical for immunopathogenesis during TB. The function of neutrophils in TB pathogenesis is believed to be via T-cell suppression by MDSC like activity and creating a permissive intracellular niche for Mtb growth. Since we did not observe any MDSC like activity of neutrophils *in vivo* (Extended Data Figure 2d) *in vitro* (Ext Data Fig. 6a), we propose that IFN-I mediated impairment of T-cell responses might be a consequence of systemic effects exerted by bacteria permissive neutrophils.

IFN-I cytokines posess potent hematopoiesis modulating functions as they cause hematopoietic stem and progenitor cell collapse to blunt monocyte production during acute infections^68^. Chronic Mtb H37Rv infection in mice promotes lymphopoiesis and suppresses myelopoiesis by altering heme metabolism in myeloid progenitors^69^. In support of these studies, Ifnar1 inhibition restored CMP frequency within bm cells (Fig.6 and 7). However, we also showed that Mtb HN878, unlike the relatively less virulent strains, preferentially expands GMPs and iGMPs by engaging Ifnar1. The overwhelming IFN-I responses induced by this strain in the susceptible *Il1r1^−/−^* and C3HeB mice promotes a granulocyte-biased hematopoiesis evidenced by a greater GMP/CLP ratio compared to the relatively resistant C57BL/6 mice, where IFN-I is tightly controlled by the sst1^R^ loci generating a balanced production of myeloid cells and lymphocytes. One likely explanation to this phenomenon is by driving a granulocyte-skewed hematopoietic response, highly virulent Mtb strains fosters neutrophil production to inflict tissue damage and survive in these nutrient-replete inflammatory consolidations^21,40^. Moreover, by inducing neutrophil production, the lineage 2 strains perhaps maintain a high G-CSF level in the host that by a feed forward mechanism provides a continuous supply of pathogen permissive neutrophils.

In summary, our study uncovered an inflammatory pathway consisting of neutrophil-GCSF-IFN-I that impairs the protective T-cell immunity in the lung and promotes immunopathogenesis during TB (Extended Data Fig.10). Although these pathways offer targets for host-directed interventions, the pleiotropic nature of these cytokines is a major bottleneck. Future studies are warranted to identify the downstream signaling targets involve in regulating pathologic granulopoiesis and neutrophil-mediated immunopathogenesis to boost protective immunity.

## METHODS

### Ethics statement

This study was approved by the Institutional Review Board of the Shenzhen University School of Medicine, China, and written informed consent was obtained from each participant. All protocols involved human samples had been approved institutional Review Board of the Shenzhen University School of Medicine.

### Human subjects and samples

Three independent cohorts were recruited from Shenzhen Third People’s Hospital (China) and Shenzhen Baoan People’s Hospital (China). Cohort I included 100 healthy controls (HC) and 103 TB patients. Cohort II included 20 HCs, 20 LTBI and 20 TB patients whom serums were obtained for G-CSF analysis. Cohort III included 226 cases of TB for evaluating NLR levels and disease status. All TB patients had no record of prior TB disease or anti-TB chemotherapy at the time of enrolment. Diagnosis of active TB was based on clinical symptoms, chest radiography and microscopy for acid fast bacilli (AFB), sputum Mtb culture. HC with normal chest radiographic findings and without a clinical history of TB were recruited. Mtb-specific interferon gamma release assays (IGRA) were used to differentiate individuals with LTBI from HC without infection as described previously^70^. The characteristics of all cohorts are summarized in supplementary figure 1. All samples from patients were collected before initiation of antibiotic treatment. ESR and WBC values are routinely obtained at the recruitment sites. Aliquots of plasma samples were used to measure G-CSF by ELISA (Cat# EHC132, Neobioscience).

### High-resolution computed tomography examination and radiological scoring

The HRCT scores were based on the percentage of lung parenchyma abnormality as previously described^71^. HRCT were performed at 10 mm section interval with a window width between 300 and 1600 Hounsfield Units and a window level between 2550 and 40 Hounsfield Units using the Toshiba Aquilion 64 CT Scanner (Toshiba, Japan). HRCT scans were analyzed two independent chest radiologists.

### Mice

C57BL/6J (Stock#00664), *Nos2^−/−^* (B6.129P2-Nos2^tm1Lau^/J, stock#002609), *Il1r1^−/−^* (B6.129S7-*Il1r1^tm1Imx^*/J, stock#003245), C3HeB/FeJ (stock#000658), B6.SJL-*Ptprc^a^ Pepc^b^*/BoyJ (stock#002014) carrying the pan leukocyte marker CD45.1 or Ly-5.1, and Ifnar1*^−/−^* (B6(Cg)-Ifnar1^tm1.2Ees^/J, stock#028288). Mice were housed under specific pathogen-free conditions. Mouse studies at the Albany Medical College (AMC) were performed using protocols approved by the AMC Institutional Animal Care and Use Committee (IACUC) (Animal Care User Protocol Number ACUP-21-04003) in a manner designed to minimize pain and suffering in Mtb-infected animals. Any animal that exhibited severe disease signs was immediately euthanized in accordance with IACUC approved endpoints.

### Mouse infection

The wild type strains of *M. tuberculosis* (Mtb) used in these studies was PDIM positive H37Rv, Mtb HN878. Mtb HN878 strain expressing msfYFP was generated by transforming the plasmid PMV261-msfYFP that constitutively expresses msfYFP under the control of the hsp60 promoter (kind gift from Dr. Christopher Sassetti, UMASS Chan Medical School, Worcester, MA). Bacteria were cultured in 7H9 medium containing 0.05% Tween 80 and OADC enrichment (Becton Dickinson). For mouse infections, mycobacteria were suspended in phosphate-buffered saline (PBS)-Tween 80 (0.05%); clumps were dissociated by sonication, and ~100 CFU were delivered via the respiratory route using an aerosol generation device (Glas-Col IES, Terre Haute, Indiana) as described previously^21^.

### Histopathology

Lung tissues were fixed in 10% buffered formalin and embedded in paraffin. Five micrometer-thick sections were stained with hematoxylin and eosin. All staining was done by the histopathology core facility at the Albany Medical College. Images of the H&E-stained lung sections were acquired by the NanoZoomer 2.0 RS Hamamatsu slide scanner. All the images were analyzed using the Image J software.

### Immunofluorescence Microscopy

Paraffin embedded lung tissue sections were cut at 5 μm thickness, mounted on ultraclean glass slides and kept for overnight drying. Tissue sections were deparaffinized and rehydrated using the following steps: xylene for 10 min, ethanol solutions (100, 95, 70, 55 and 30 % for 5 min each), and distilled water for 20 min. Slides were subjected to antigen retrieval by boiling in Tris-EDTA buffer (pH=9.0) for 15 min. Slides were transferred in distilled water to cool them to room temperature. For permeabilization, sections were dipped in 0.4 % Tween 20 in TBS (TBST) for 30 min. The sections were washed twice for 5 min with wash buffer (0.05 % TBST). Nonspecific antibody binding sites were blocked by incubating tissue sections with 5 % BSA for 2 hrs at room temperature. Sections were rinsed in wash buffer and incubated overnight in primary antibodies against the Ly6G, CSF3, or CSF3R. As controls, pre-immune serum and isotype matched controls were used. After incubation, the tissues were washed 3 times for 5 minutes with wash buffer at room temperature and incubated in the respective secondary antibodies (anti-rabbit conjugated to Alexa-488, anti-rat conjugated to Alexa 594) for at least 2h at room temperature. After secondary antibody incubation, tissue sections were washed several times each for 5 minutes. All secondary antibodies used for the immunostaining were cross absorbed. Tissue sections were mounted using Prolong Gold Antifade reagent (Invitrogen, grand Island, NY) with DAPI, and the tissue sections were examined in ECHO Revolve 4 microscope. Isotype matched control antibodies were used for checking antibody specificity.

### Treatment of animals with antibodies or cytokines

#### rmG-CSF

Recombinant murine G-CSF (100ug/kg,) was administered to naiive mice every day for 5 consecutive days via the intraperitoneal route (i.p.). The next day, animals were euthanized, organs were analyzed by flow cytometry for the presence of immune cells or progenitor cells. In Mtb HN878 infected mice, recombinant murine G-CSF (100ug/kg) was administered to mice that were either treated with anti-IFNAR1 blocking antibody or the IgG control antibody starting at day 19, every other day till day 27. Animals were euthanized on day 28 to analyze bacterial burden, immune cell infiltration and bone marrow progenitors.

#### G-CSF blockade

Anti-G-CSF neutralizing antibody (2.5mg/kg, i.p.) was administered to mice starting at 11 days post infection (dpi), every alternate day till day 27. Control cohorts received IgG2 Isotype control antibody. Mice were euthanized on day 28, and organs were collected for flow cytometry and bacterial burden analysis by CFU assay.

#### IFNAR1 blockade

Anti-IFNAR1 neutralizing antibody was administered to mice (25mg/kg, i.p.) at alternative days from 9-27 dpi. Control cohorts received IgG2 Isotype control antibody. Mice were euthanized on day 28 and organs were collected for flow cytometry and CFU analysis.

#### Neutrophil depletion

Anti-Ly-6G depleting antibody (250 ug/mouse) or isotype control antibody (2A3) was administered to animals via i.p. route every alternative day from 11-27 dpi. Ly-6G^+^ neutrophil depletion was verified by FACS analysis of CD11b^+^ Gr-1^+^ cell populations and animals were euthanized on 28 dpi to determine bacterial burden by CFU, histopathology by hematoxylin and eosin staining, cytokines in the lung by bioplex, and quantify the progenitors in the bone marrow.

### Flow Cytometry

Tissues were collected in ice-cold PBS and subjected to preparation of single cell suspensions. Lung tissues were digested with Collagenase type IV (150U/mL)/DNase I (60U/mL) and passed through 40 μm cell strainers to obtain single cell suspension. Bone marrow cells were harvested by flushing femur bone with 10 ml 1X PBS (pH 7.0 ±0.2) while the spleen tissues were crushed through 40 μm cell strainers using FACS buffer (1X PBS+0.5% BSA). The single cell suspensions were subjected to erythrocyte lysis using ACK lysis buffer and non-specific antibody binding sites were blocked by Fc-Block CD16/32. The cells were surface stained with directly conjugated antibodies (Table) for 30 min at 4 °C in dark in FACs buffer. Subsequently, cells were washed with FACs buffer and fixed for 20 minutes at 4 ^°^C using the fixation buffer. For intracellular staining, surface-stained cells were washed with 1X Perm-wash buffer and fixed with Cytofix-cytoperm for 20 min at 4 °C. Subsequently, the fixed cells were washed with Perm-wash buffer and stained with directly conjugated antibodies (Table) in FACs buffer for 30 minutes at 4 °C in dark. For Ki67 staining, cells were fixed and permeabilized with buffer from foxp-3 intracellular staining kit, prior staining. For analyzing granulocyte macrophage progenitors (GMP), bone marrow cells were first stained with an anti-mouse lineage antibody cocktail, anti-mouse c-Kit-APC, anti-mouse Sca1-PercpCy5.5, anti-mouse CD34-PE, anti-mouse CD16/32-PE cy7 and anti-mouse CD127-Alexa 488. Specifically, for GMP analysis, Fc-block (CD16/32) antibody was not used before surface staining. Events were acquired in LSRII/BD Symphony flow cytometer and analyzed using FlowJo v.10. All analyses were performed on live cells by excluding dead cells by using the fixable viability stain conjugated with eFlour780. Gating strategies to identify specific cell populations are provided in applicable figures.

### Cytokine measurement and immunoblotting in tissue homogenates

Murine cytokine/chemokine concentrations in lung homogenates from Mtb infected mice were quantified using the Bio-Plex Pro^TM^ Mouse Cytokine Plex assay (Table x). Briefly, lung tissues were collected from mice on mentioned days, flushed with sterile 1X PBS (pH 7.0±0.2) and homogenized in ice cold PBS. The concentration of total protein in the lung homogenates were quantified using the Pierce BCA protein assay kit according to the manufacturer’s instructions. The BioPlex assay was performed according to manufacturer’s instructions and fluorescence was detected using a Bioplex-200 reader. Data analysis was performed with Biolpex Data Pro Software and concentration of cytokine/chemokine in pg/mg total protein was calculated and depicted in the graphs.

### Mixed bone marrow chimera generation

*Il1r1^−/−^* mice were lethally irradiated with two doses of 600 rads with a 3hr interval between 1^st^ and second dose. After the 2^nd^ irradiation, bone marrow from CD45.1^+^ wild type mice (*Il1r1^+/+^*) and CD45.2^+^ knockout mice (*Ifnar1^−/−^*) was isolated, red blood cells were lysed using ACK lysis buffer (Lonza, Cat# BP10-548E) and the remaining cells were quantified using an automated cell counter (Bio-Rad). CD45.1+ and CD45.2+ cells were then mixed equally at a 1:1 ratio and 10^7^ cells from this mixture were injected intravenously into lethally irradiated hosts. Reconstitution of host mice lasted 6-8 weeks with preventative sulfatrim treatment for the first 4 weeks. Mixed bone marrow chimera mice were then infected by low dose aerosol with *M. tuberculosis* HN878 as described previously. 4 weeks following infection, the lungs of chimera mice were harvested, and single cell suspensions were made following collagenase (150U/ml) and DNaseI (60U/ml) treatment and tissue disruption. Single cell suspension of lung was further used for flow cytometry analysis of neutrophils, T-cells (CD4/CD8) among lung leukocytes. The femurs from the mixed chimera mice were harvested, bone marrow (BM) cells were prepared and stained for myeloid and lymphoid progenitors. Briefly, BM cells were stained with lineage antibody cocktail (Biolegend) followed by c-Kit, Sca1, CD16/32 and CD127 antibodies to identify myeloid and lymphoid progenitors as shown in Extended Data Fig. 4. The details of the clone and catalog number of the antibodies were given in Supplementary table 3.

### Isolation of Neutrophils from Mtb infected mice

Single cell suspension of the lung, spleen and bone marrow were prepared from C57BL/6J and *Il1r1^−/−^* mice, red blood cells were lysed using ACK lysis buffer, and filtered through 40um filter. Mouse neutrophils were prepared by negative selection using the Mojosort neutrophil isolation kit from Biolegend (Cat#480058) according to the manufacturer’s protocol. Subsequently, positive selection of the neutrophils was performed by staining the flow through cells with anti-Ly-6G-APC for 20 minutes at 4^0^C followed by isolating the stained cells with anti-APC nanobeads using a mojosort APC nanobead kit (Biolegend). Purity and viability of the cells were checked by flow cytometry using an anti-mouse Gr1 antibody and fixable viability dye. Up to 90-95% purity was achieved using this cell isolation protocol.

### RNA Sequencing

Ly6G+ cells were isolated from *Mtb* infected C57BL/6 and *Il1r1^−/−^* mouse lungs (n=4/strain), spleen(n=4/strain) and bone marrow (n=4/strain) using the Mojosort neutrophil isolation kit and Mojosort anti-APC kits as described previously, following the manufacturer’s instructions. All RNA samples were subjected to rigorous quality control and samples with a RIN>6.0 were chosen for further processing. RNA sequencing libraries were prepared using the NEBNext Ultra II RNA Library Prep Kit for Illumina following manufacturer’s instructions (NEB, Ipswich, MA, USA). Briefly, mRNAs were first enriched with Oligo(dT) beads. Enriched mRNAs were fragmented for 15 minutes at 94 °C. First strand and second strand cDNAs were subsequently synthesized. cDNA fragments were end repaired and adenylated at 3’ends, and universal adapters were ligated to cDNA fragments, followed by index addition and library enrichment by limited-cycle PCR. The sequencing libraries were validated on the Agilent TapeStation (Agilent Technologies, Palo Alto, CA, USA), and quantified by using Qubit 2.0 Fluorometer (Invitrogen, Carlsbad, CA) as well as by quantitative PCR (KAPA Biosystems, Wilmington, MA, USA). The sequencing libraries were clustered onto 2 flowcell lanes. After clustering, the flowcell were loaded on the Illumina HiSeq instrument (4000 or equivalent) according to manufacturer’s instructions. The samples were sequenced using a 2×150bp Paired End (PE) configuration. Image analysis and base calling were conducted by the HiSeq Control Software (HCS). Raw sequence data (.bcl files) generated from Illumina HiSeq was converted into fastq files and de-multiplexed using Illumina’s bcl2fastq 2.17 software. One mismatch was allowed for index sequence identification.

### RNAseq Data Analysis

After investigating the quality of the raw data, sequence reads were trimmed to remove possible adapter sequences and nucleotides with poor quality using Trimmomatic v.0.36. The trimmed reads were mapped to the Mus musculus reference genome available on ENSEMBL using the STAR aligner v.2.5.2b. The STAR aligner is a splice aligner that detects splice junctions and incorporates them to help align the entire read sequences. BAM files were generated as a result of this step. Unique gene hit counts were calculated by using feature Counts from the Subread package v.1.5.2. Only unique reads that fall within exon regions were counted. After extraction of gene hit counts, the gene hit counts table was used for downstream differential expression analysis. Using DESeq2, a comparison of gene expression between the groups of samples was performed. The Wald test was used to generate p-values and Log2 fold changes. Genes with adjusted p-values < 0.05 and absolute log2 fold changes > 1 were called as differentially expressed genes for each comparison. A gene ontology analysis was performed on the statistically significant set of genes by implementing the software GeneSCF. The GOA_human or mouse GO list was used to cluster the set of genes based on their biological process and determine their statistical significance. A PCA analysis was performed using the “plot PCA” function within the DESeq2 R package. The plot shows the samples in a 2D plane spanned by their first two principal components. The top 500 genes, selected by highest row variance, were used to generate the plot. RNAseq raw fastq reads were mapped to mouse reference using STAR and the raw read counts for each gene were calculated using HTseq with human gene annotation GTF file downloaded from gene code (https://www.gencodegenes.org/human/). RNAseq data were subsequently normalized based on the negative binomial distribution, with variance and mean linked by local regression, and differentially expressed genes among different sample groups, including spleen (BS vs. RS), bone marrow (BM vs. RBM) and lung (BL vs. RL), were determined using the Bioinformatics Toolbox of MATLAB (version 2021a) according to (https://www.mathworks.com/help/bioinfo/ug/identifying-differentially-expressed-genes-from-rna-seq-data.html).

### (A) Manually curated gene set for enrichment analysis

Previously published TB gene expression modules based on WGCNA analysis for human and mouse TB microarray expression data were obtained from Moreira-Teixeira et al^24^ and Singhania et al^18^. There were 23 human and 27 mouse TB modules. In terms of human TB modules, all genes involved in these 23 human TB modules were provided by Moreira-Teixeira et al^24^. These human genes were mapped to its murine homologs based on the lookup table of human and MGI (http://www.informatics.jax.org/downloads/reports/HOM_MouseHumanSequence.rpt)^72,73^ and were further used for gene set enrichment analysis with GSVA^74^.

### (B) Gene set enrichment analysis and differential expression analyses

Enrichment of the TB gene signatures and neutrophils related human gene sets was performed according to Moreira-Teixeira et al^24^ using the R package GSVA^74^. There were 23 human and 27 mouse gene sets for the TB gene signatures, while 145 neutrophils related human gene sets were extracted from the curated GSEA C2 gene sets downloaded from GSEA website. In details, we carried out on a per sample basis gene set enrichment analysis using ssGSEA in GSVA. Compared to traditional gene set enrichment analysis, such as GSEA^75,76^, the enrichment scores from GSVA were based on absolute expression rather than differential expression to quantify the degree to which a gene set is over-represented in a particular sample. As each sample being determined for gene set enrichment, we used GSVA enrichment scores to determine whether a gene set was significantly differently enriched among three TB groups, including BS vs. RS, BM vs. RBM and BL vs. RL. We prioritized gene sets showing significant difference (t-test P<0.05 conducted by muttest in MATLAT 2021a) in the comparison of BL vs. RL. Furthermore, we also required that the gene set was significant either in the comparison of BM vs. RBM or BS vs. RS. Gene sets fulfilled the above criteria were further followed by differential expression analysis for genes included in these TB modules. Similarly, we performed differential expression analysis using the same t-test statistic method for these selected genes emerged from the above procedure. Only genes showing the significantly differential expression among BL vs. RL and either in the comparison of BM vs. RBM or BS vs. RS, were subjected to heatmap visualization. We further selected these top genes from these enriched TB modules using the absolute fold change >2.

### (C) IPA analysis

Predictive pathway analysis was performed using Qiagen® Ingenuity Pathway Analysis software (Qiagen) which compared significantly different–log_2_ gene expression values between test groups.

### Cell DIVE Multiplexed immunofluorescence

Multiplex experiments were conducted on a total of 6 biospecimens using Cell DIVE™, a hyperplexed imaging technique that allows for spatial biomarker mapping for more than 60 biomarkers using an iterative staining, imaging, and dye inactivation workflow (Figure 1j). Multiplexed immunofluorescence (MxIF) imaging was performed at GE Research using methodologies described in detail in Gerdes et. al^32^. In brief, eleven antibodies (Supplementary Table 3) were stained after deparaffinization, rehydration, and a single two-step antigen retrieval process. Antigens were detected using either primary antibodies with fluorophore-labelled secondary antibodies or fluorescent dye conjugated antibodies. Prior to antibody staining, tissues were stained with DAPI and imaged in all channels of interest to collect inherent tissue autofluorescence (AF). Whole slide images at 20x were collected each round using DAPI, FITC, Cy3, and Cy5 channels on an IN Cell 2200, followed by automated AF subtraction, registration with baseline DAPI, and region stitching. Pseudo-colored virtual H&E images were also generated using DAPI and AF signal. Tissue was stained with 2-3 antibodies per cycle, imaged, and then dye inactivated to remove signal. The MxIF cycling was repeated until all target antigens were probed. Biomarkers included *Mycobacterium tuberculosis*-associated proteins (Mtb) and immunological (B220, CD3e, F4/80, Ly-6G), and cell proliferation marker Ki67. The typical antibody characterization workflow is a standardized process that consists of screening at least three commercial antibody clones per target for sensitivity and selectivity, evaluation of the epitope for susceptibility to the dye inactivation workflow, and concordance of antibody performance upon dye conjugation (Details of the antibodies and clones used for Cell Dive was listed in Supplementary Table 3).

### Single Cell Analysis of TB lesions using QuPath

Percent population of immunostained cells were quantitatively assessed within regions of interest (ROIs) using open source QuPath 0.2.3 digital software. Whole slide images (WSI) of lung lobes were annotated for inflamed (n=3) and mildly inflammation/uninvolved (n=3) ROIs using the rectangle tool, defined by visual scoring of *Mycobacterium tuberculosis* immunoexpression (Invitrogen PA1-73136) as shown in Extended Data Fig 1g. Single cell segmentation was run using QuPath’s inbuilt cell detection command (Extended Data Fig 1g) using the DAPI (nuclei) channel and expansion of the cytoplasm to a predefined distance. Subclassification of each segmented object was performed using a pixel-based object classifier to identify cell populations of interest (B220+, CD3+, F4/80+, Ki67+, Ly6G+, Mtb+). The measurement map tool was employed to visualize biomarker staining intensity across all annotations and threshold parameters for each object classifier were constrained across all samples and ROIs. Secondary composite classification (Extended Data Fig 1g) was performed to classify Ki67+CD3+, Ly6G+Mtb+, and F4/80+Mtb+ populations. The number of positive cells present were calculated as a proportion of the total number of cells detected in each ROI.

## Statistical analysis

The statistical significance of differences between data groups was determined using the Unpaired two tailed student’s *t*-test, or one-way ANOVA using Tukey’s multiple comparison test using Graphpad Prism 9 (San Diego, CA); * (p<0.05) ** (p<0.001) *** (p < 0.0001) denotes significant differences compared as indicated in figures. Equal variance between samples was assessed by Brown-Forsythe test. Experiments in which variances were equivalent, were analyzed by one-way ANOVA with Sidak’s multiple comparisons test. Those with unequal variances were analyzed by Welch ANOVA and Dunnett’s multiple comparisons test.” All survival data are presented as Kaplan-Meier plots.

## Data Availability

The RNA-seq datasets of mouse neutrophils have been deposited in the NCBI Gene Expression Omnibus database with the accession number GSE203118. Publicly available datasets used in this study include GSE19419, GSE94438, GSE144127, GSE112104 and GSE79362.

## Author contributions

Conceptualization and design: BBM and MS. Human study: YC, XC, YY, QY, FL, WW. Experiments: MS, BBM, EM, PS. Data analysis and interpretation: BBM, MS, EM, YC. Bioinformatics: ZZ and SD. Reporter strain generation: AKO. Cell Dive imaging and data analysis: EM, FG and BBM. Manuscript writing: BBM, MS. BBM and YC designed the human study and YC oversaw the all the studies involving human subjects and samples. BBM oversaw the entire study and acquired funding.

## Supporting information

Information of human cohorts used in the study

Differentially expressed genes in neutrophils isolated from lung, spleen and bone marrow of wild type and Il1r1 KO mice

Reagents and resources used in the study

## Acknowledgement

This study is supported by NIH grant (AI148239-01A1 to BBM) and National Natural Science Foundation of China (grant 82072252 to YC).

## FIGURE LEGENDS

**Extended Data Fig. 1. Higher neutrophil to T-lymphocyte ratio predicts disease outcome, related to Fig. 1**. Indicated mouse strains were infected by aerosol challenge of Mtb HN878. At 28 dpi, mice were sacrificed, and various disease parameters were evaluated. (**a**) Representative images of H&E-stained lung sections for the indicated strains. (**b**) Lung bacterial burden was presented as CFU. (**c**) Influx of neutrophils and (d) CD3e^+^ T-cells in the single cell suspensions prepared from the lung of indicated strains determined by flow cytometry. (**e**) Relative abundance of neutrophils to CD3e^+^ T-cells in the lung. (**f**) the Number of neutrophils and (g) NLR in AFB- or AFB+ TB patients. (**h**) Correlation of the HRCT score (left) and ESR (right) with NLR in TB patients, N=226. (**i**) Flow cytometry quantitation of infected myeloid cells, representative fluorescent images of cytospins prepared from the wt and *Il1r1^−/−^* mice lung. (**j**) Schematic representation of Cell DIVE multiplexed immunofluorescence and image analysis, related to **Fig 1j**. Three regions of interest (ROI) each were chosen randomly based on the magnitude of “Ly6G intensity” and “presence of bacteria” (defined as “inflamed”, yellow boxes); and with relatively less intensity for Ly6G and Mtb or completely devoid of noticeable inflammation uninvolved/uninfected regions (defined as “mildly inflamed/uninvolved”, green boxes).

**Extended Data Fig. 2. Anti-Ly6G treatment specifically depleted neutrophils, related to fig 2**. Neutrophil depletion was carried out in the *Il1r1^−/−^* mice as shown in Fig. 2a. (a) Representative FACs plots showing depletion of neutrophils (CD11b^+^Gr1^hi^ gate shown) in the lung, spleen, and bone marrow of the indicated groups. (**b**) Effect of anti-Ly6G administration on lung cells was quantified in the total single cells prepared from the lung. (**c**) Number of T-cells expressing Ki67 in lung and (**d**) spleen related to Fig 2d-e. (**e**) Representative FACS plots showing IL-2 expressing CD4 and CD8 T-cells in the lung and spleen of indicated mouse strains. Sample size of N=6 mice were included for each group. Data are presented as Mean±SD, ***; p<0.001, Two-way ANOVA with Bonferroni’s multiple comparison-test. ns: non-significant (p-value>0.05). The data are representative of two individual experiments.

**Extended Data Fig. 3. Gating strategy for identifying hematopoietic progenitors in the bone marrow**. *Il1r1^−/−^* mice were infected with ~100 CFU by aerosol challenge of Mtb HN878 and 4 wks post infection (wpi), bone marrow single cell suspension was prepared. Cells were gated for FSC-A against SSC-A. Doublets were excluded using FSC-H against FSC-A. Viable cells were gated and lineage-committed cells were excluded. The lineage^−ve^ population was further gated for cKit expression. Lin^−^ckit^+^ cells were further divided into Sca1^−/low^ cells and Sca1^+/hi^ ^cells^. Lin^−^cKit^+^Sca1^−/low^ cells were gated on CD127^+^ and CD127^−^ cells. Lin^−^cKit^+^Sca1^−/low^CD127^+^ cells were defined as CLP. Lin^−^cKit^+^Sca1^−/low^CD127^−^ cells were further gated based on CD34 and CD16/32 to define CMP, GMP and MEP representing the “canonical hematopoiesis” response. In another strategy, the lin^−^cKit^+^Sca1^+/hi^ cells were gated based on CD127 expression and lin^−^cKit^+^Sca1^+/hi^CD127^+^ cells were defined as iCLPs. The lin^−^ckit^+^Sca1^+/hi^CD127^−^ cells were gated based on CD34 and CD16/32 to define iCMPs, iGMPs and iMEPs representing “infection-induced hematopoiesis” response.

**Extended Data Fig. 4. Hematopoiesis is regulated by bacterial virulence**. C3HeB/FeJ mice were infected with ~100 CFU of aerosolized wild type (H37Rv) or esx1 deficient (Δesx1) Mtb strains. 4 wpi, animals were sacrificed, and lineage committed progenitors within the lineage^−ve^ bone marrow cells depicting (**a**) canonical hematopoiesis and (**b**) infection induced hematopoiesis was quantified as mentioned in Extended Data Fig. 3. (**c**) Indicated strains of mice were infected with Mtb HN878 or Mtb H37Rv or Mtb Δesx1. At 4 wpi, animals were sacrificed, and relative abundance of GMP to CLP were evaluated. Sample size of N=5 was included for each group. Data are presented as Mean±SD, *; p<0.05; **, p<0.01. two-way (a-b), one-way ANOVA with Bonferroni multiple comparison test (c-d). The data shown are from one experiment.

**Extended Data Fig. 5. G-CSF inhibition potentiates T-cell functions, related to Figure 3.** (**a**) *Il1r1^−/−^* mice were infected with Mtb HN878 and were treated with anti-mouse G-CSF or isotype (n=4 mice each) as shown in the schematics and mentioned in Fig. 3d. (a) Effect of G-CSF inhibition on myeloid cells of the lung. (**b**) Neutrophil number, (**c**) number and percentage of T (CD4 and CD8) cells expressing Ki67 in the spleen determined by flow cytometry. (**d**) Representative FACS plots showing T-bet expression in CD4 and CD8 T-cells from spleen in indicated cohorts. (**e**) Representative histograms showing cytokine expressing CD4 and CD8 T-cells in the lung (upper panel) and spleen (lower panel). (**f**) T cell number expressing Ki67 or IL-2 in the spleen of C3HeB mice, infected with Mtb HN878 and treated with anti-G-CSF or isotype as shown in the schematics and mentioned in Fig. Sample size of n=6 mice were included for each group. The data are representative of two separate experiments. (**g**) C3HeB mice were infected with Mtb HN878 and treated with anti-G-CSF or anti-CSF3R as shown in the schematics. At 28 dpi mice were sacrificed, single cell suspensions from lung were prepared and analyzed for (**h**) neutrophil influx and T-cell proliferation (Ki-67^+^), (**i**) IL-2 secretion in CD4 and (**j**) CD8 T-cells by intracellular staining in the indicated cohorts. Sample size of n=5 mice were included for each group. Data shown are from one experiment. Data are presented as Mean±SD, *, p<0.05; **, p<0.01; ***; p<0.001, one-way ANOVA with Tukey’s multiple comparison-test. ns: non-significant (p-value>0.05).

**Extended Data Fig. 6. Recombinant G-CSF administration into naiive mice negatively impacted T-cell numbers and proliferation by inducing granulopoiesis and suppressing lymphopoiesis**. (**a**) Effect of recombinant G-CSF stimulation on T-cell proliferation in a neutrophil and T-cell coculture model. T-cell proliferation was measured by counting the total live CD3e^+^Ki67^+^ cells. (**b**) C3HeB/FeJ mice were treated with recombinant murine G-CSF (rG-CSF) every day for 5 consecutive days as shown in the schematics. The following day, mice were sacrificed, (**c**) Frequency of lineage-committed progenitors was calculated as described in Ext Data Fig 3. (**d**) infiltration of the number of neutrophils were quantified in the indicated organs, (**e**) number and % proliferation of CD4 and CD8 T-cells (Ki-67^+^) in the lung, (**f**) spleen were determined by flow cytometry. Each symbol represents a single mouse. PBS treated mice were used as vehicle control to assess the effect of rG-CSF on the indicated progenitors. Sample size of n=4 was included for each group. Data are presented as Mean±SD, *, p<0.05; **, p<0.01; ***, p<0.001; two-way ANOVA with Bonferroni multiple comparison test. The data shown are representative of two experiments. (**g**) C3HeB mice were infected with Mtb, and G-CSF neutralization was carried out as shown in schematics. BM progenitors were quantified, GMP/CLP ratio depicting canonical, and infection induced hematopoiesis were compared between treatment groups and shown. (**h**) Gating strategy to identify maturation stages of neutrophils in the bone marrow cells based on CD11b and Ly6G expression. CD11b^+^Ly6G^hi^, CD11b^+^Ly6G^mid^ and CD11b^+^Ly6G^−ve^ cells were defined as mature neutrophils, metamyelocytes and myelocytes respectively. CD11b^−^Ly6G^−^ cells were further gated based on CD34 and ckit expression. cKit^+^CD34^+/hi^ and cKit^+/low^CD34^+^ cells were defined as promyelocyte and myeloblasts. Representative FACs plots showing the gating strategy and table depicting the expression of various surface markers were shown. (**i**) Quantitation of different developmental stages in the neutrophil ontogeny was shown. Data are presented as Mean±SD, *; p<0.05, two-way ANOVA with Bonferroni multiple comparison test. The data are representative of two individual experiments.

**Extended Data Fig. 7. Type I IFN specific genes are enriched in neutrophils during Mtb infection, related to Fig 4**. RNA sequencing analysis of neutrophils from indicated organs of *Il1r1^−/−^* mice (R) compared to *Il1r1^+/+^*mice. (a) Volcano plots showing differentially expressed genes (DEGs) in different organs. (**b**) WGCNA analysis dot plot showing the enrichment of gene modules previously implicated in human TB. Size and color of the dot represents the number of genes and enrichment within the modules. Boxes indicate the enrichment of interferon modules in tissues. (**c**) Dot plots showing the enrichment of gene sets implicated in the indicated pathways obtained by comparing them with previously reported neutrophil specific pathways in the MsigDB database.

**Extended Data Fig. 8. Bulk RNA sequencing of neutrophils from lung, spleen and bone marrow identifies type I IFN signature and tissue specific response, related to Fig 4**. (**a**) Bi-clustering heatmap visualizing the expression profile of DEGs, as sorted by adjusted p-value via plotting the log_2_ transformed expression values in each sample. (**b**) GSEA showing enrichment of genes implicated in Ifn-α response. (**c**) Ingenuity pathway analysis of DEGs predicting activation or inhibition of upstream regulators in relation to interferon and iNos-signaling and (**d**) IL-7 signaling, cholesterol biosynthetic pathways etc. in the lung, spleen, and bone marrow of *Il1r1^−/−^* mice (R) compared to wt *Il1r1^+/+^* mice (B).

**Extended Data Fig. 9. IFNAR1 is necessary and sufficient to promote G-CSF mediated pathologic granulopoiesis.** C3HeB/FeJ mice were infected with Mtb HN878 and treated with anti-IFNAR1 or isotype (n=10 mice each). At 19 dpi, both these cohorts were administered recombinant murine G-CSF every alternate day till 27 dpi as shown (**a**) in the schematics. At 28, dpi mice were sacrificed, (**b**) Frequency of Lin^−^ckit^+^sca1^−/low^ or Lin-ckit^+^Sca1^+/hi^ lineage committed progenitors within the bone marrow cells, relative abundance of GMP verses CLP depicting conventional and (**c**) infection-induced hematopoiesis was shown. (**d**) Quantitation of different developmental stages in neutrophil ontogeny within the bone marrow of indicated cohorts. (**e**) Representative histopathological images from FFPE lung sections, stained with hematoxylin and eosin (**f**) Absolute number of neutrophils, (**g**) infected neutrophils, (**h**) CD4 and CD8 T-cells and (**i**) relative abundance of neutrophils versus CD4 or CD8 T-cells were determined in the lungs of these cohorts. Data are presented as Mean±SD, Statistical significance was calculated by two-way ANOVA with Bonferroni multiple comparison test. *, p<0.05; **; p<0.01; ***, p<0.001. Data shown are from two independent experiments with 5 mice per group. Source data are provided as a Source data file.

**Extended Data Fig. 10. Model to describe the role of G-CSF/IFN-I in impairing T-cell immunity and driving immunopathogenesis during TB**. Mtb infection induces robust neutrophil infiltration to the lung. These cells secrete G-CSF to promote granulocyte-biased hematopoiesis in the BM characterized by an expansion of GMPs relative to CLPs that compromises lymphocyte production. The preferential expansion of GMPs results in rapid production and mobilization of both immature and mature neutrophils to the site of infection in an IFNAR1-dependent fashion. The IFNAR1 expression on neutrophils is critical for their recruitment to the lung and causing immunopathology. Inhibiting IFNAR1 signaling restores hematopoietic equilibrium by reducing the GMP/CLP ratio, and spatiotemporal positioning of proliferating and cytokine secreting CD4 and CD8 T-cells in the TB lesions eventually leading to containment of bacterial replication and immunopathology. This model predicts that dysregulated G-CSF/IFN-I pathways disrupt the normal hematopoietic response in the bone marrow that compromises T-cell response in TB lesions and promotes pathogen replication by a continual supply of bacteria permissive neutrophils.

