## Supplementary material for "Neutrophils reprograms the bone marrow to impair T-cell immunity during tuberculosis": Supplemntary Figures

### Extended Data Figure 1

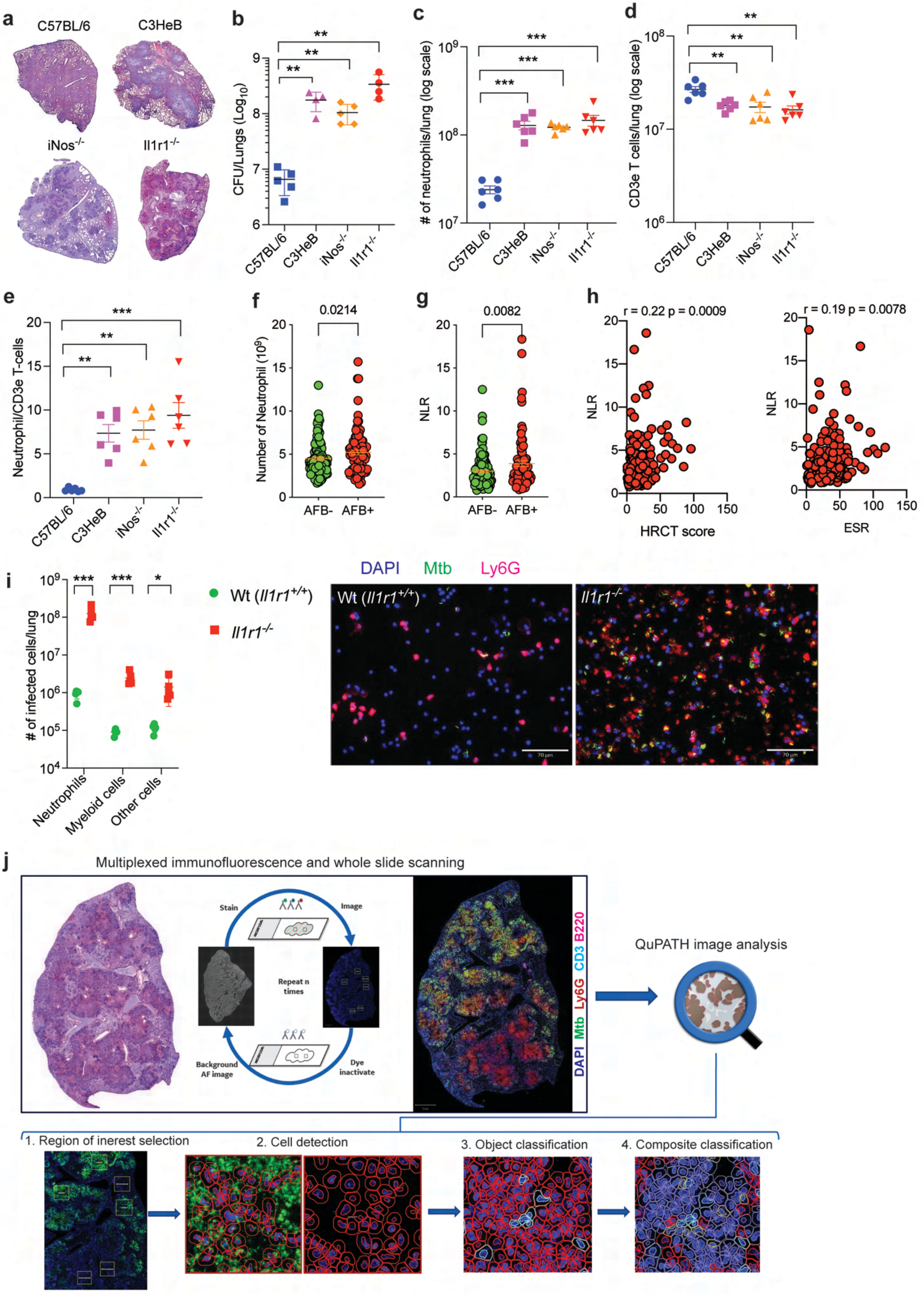

#### Extended Data Figure 2

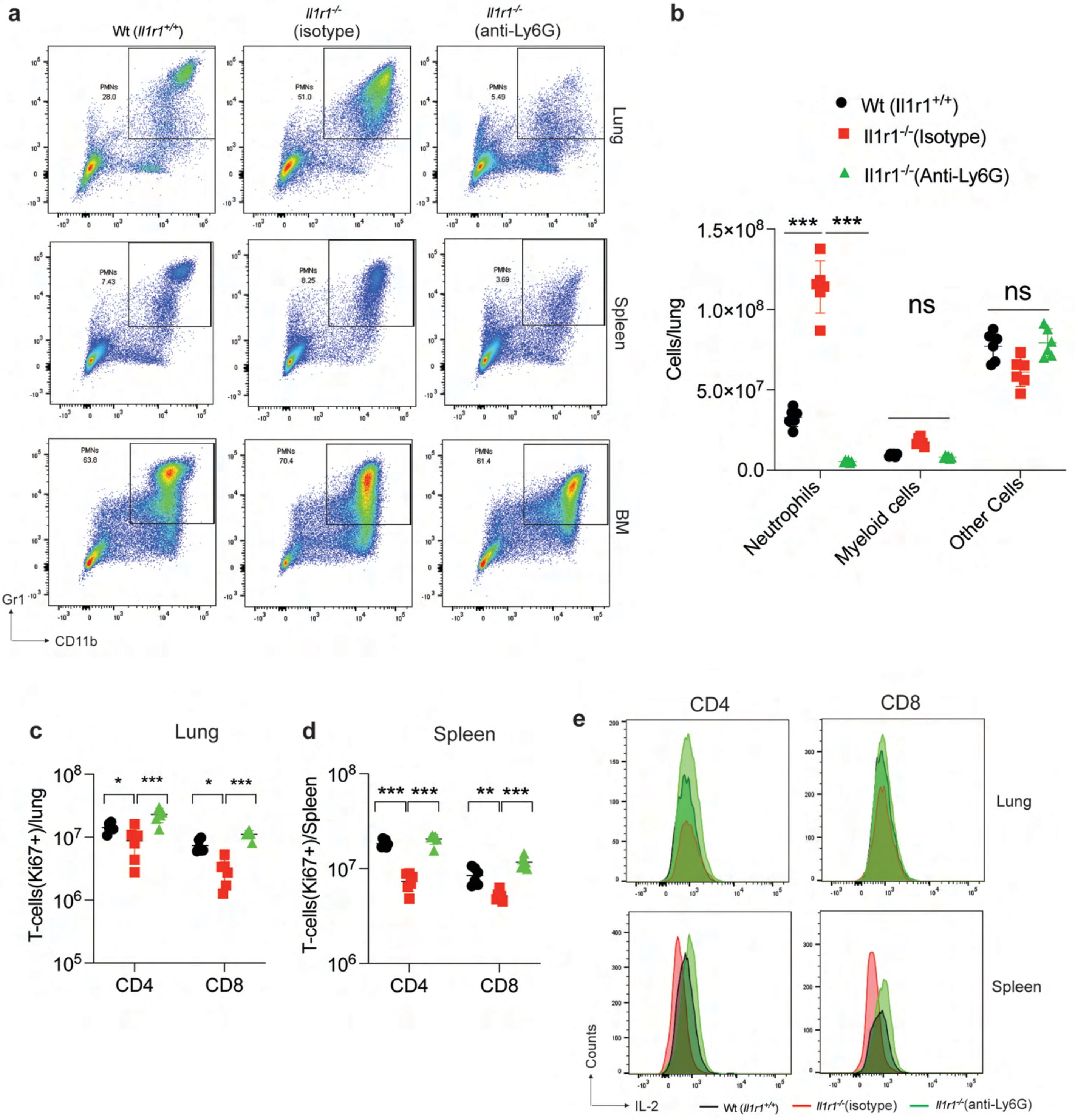

#### Extended Data Figure 3

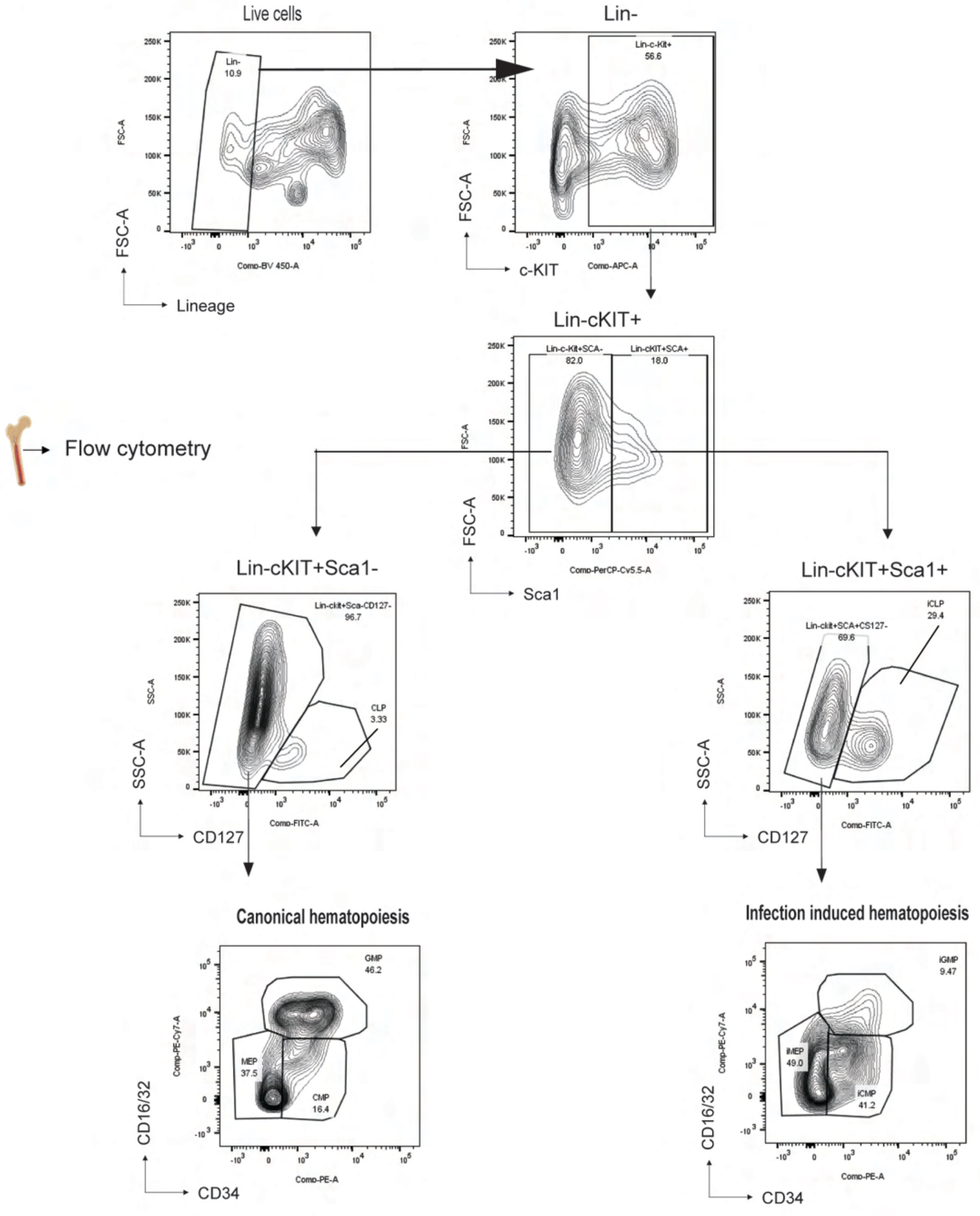

**Extended Data Figure 4**

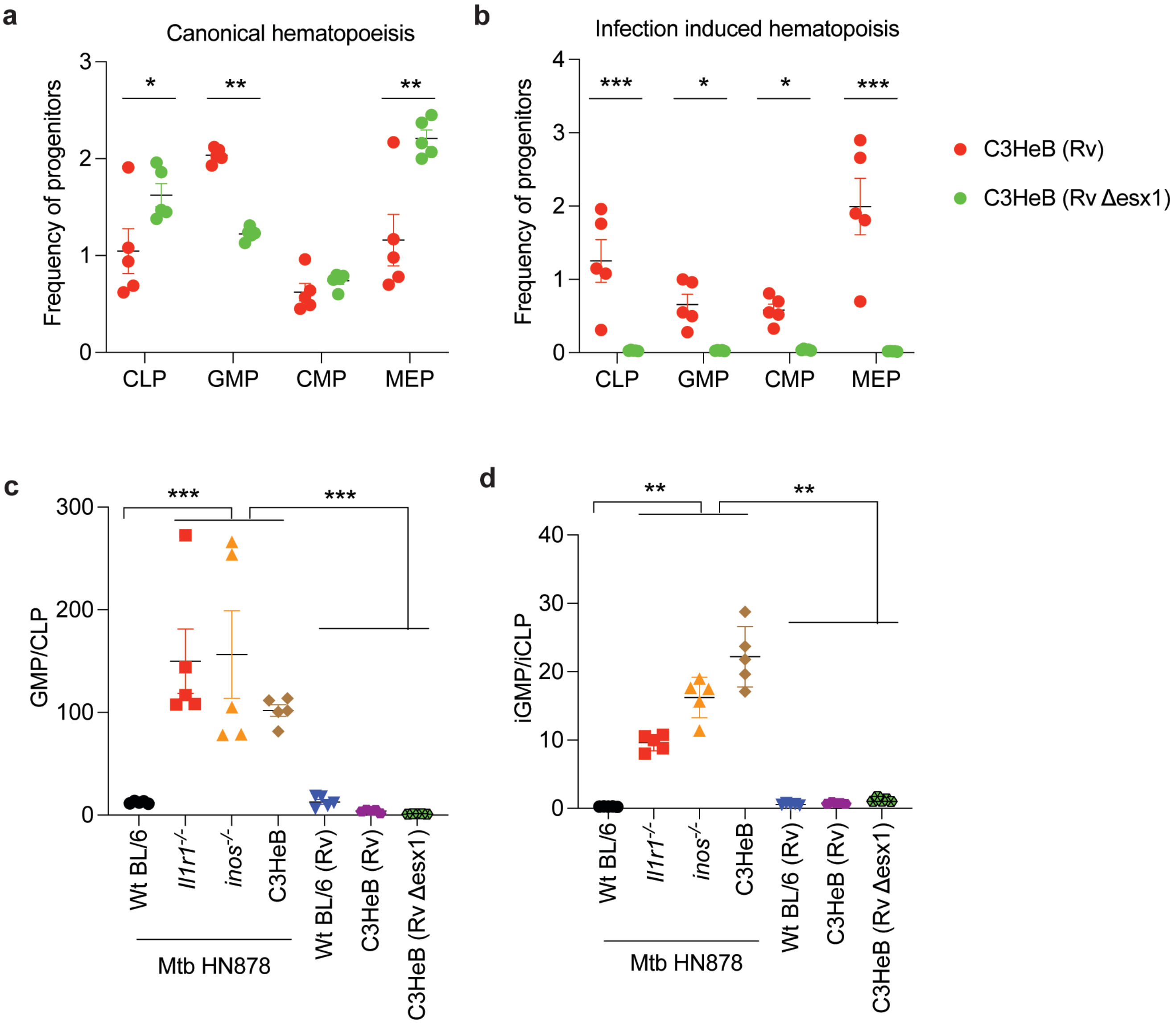

#### Extended Data Figure 5

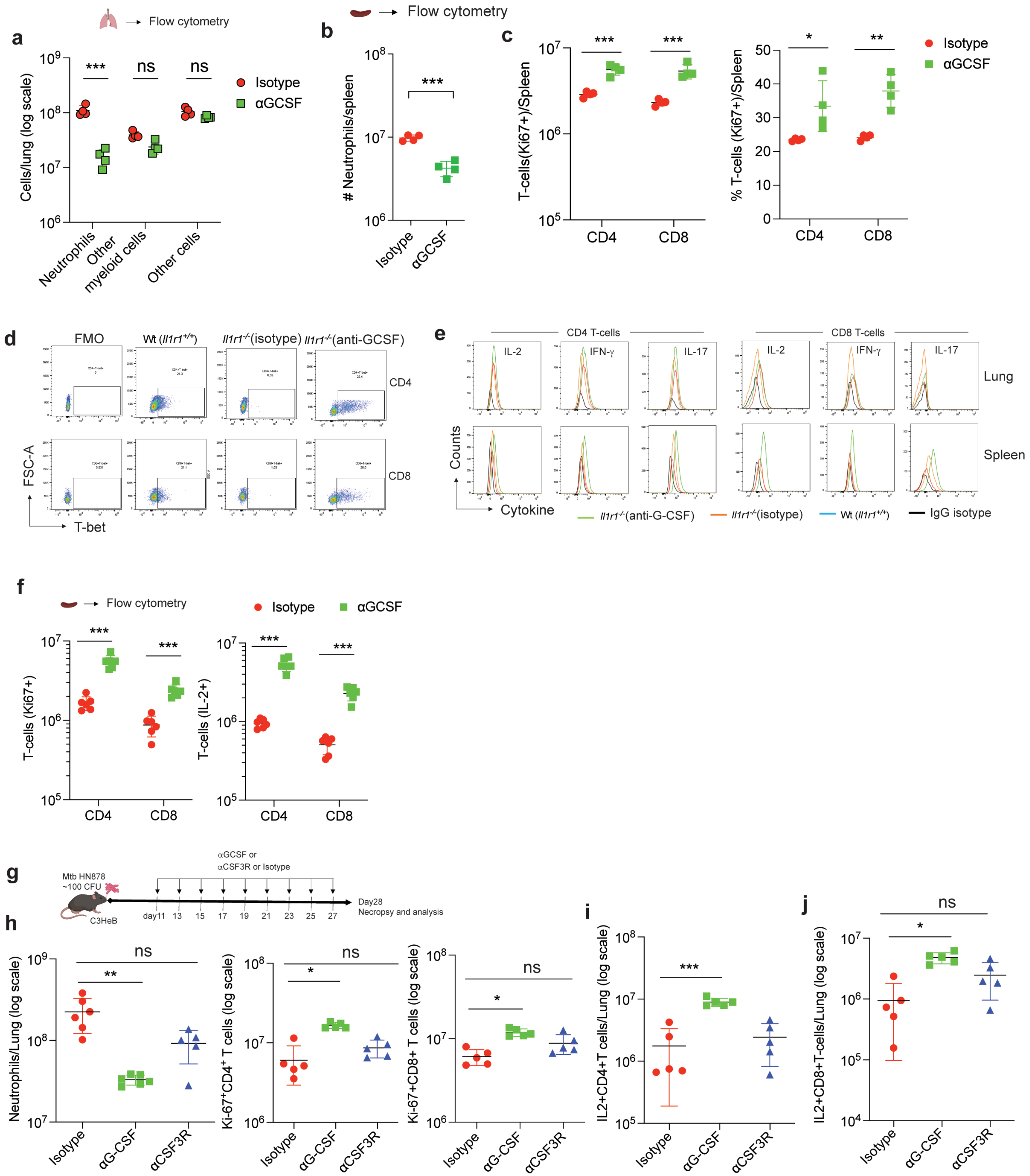

### Extended Data Figure 6

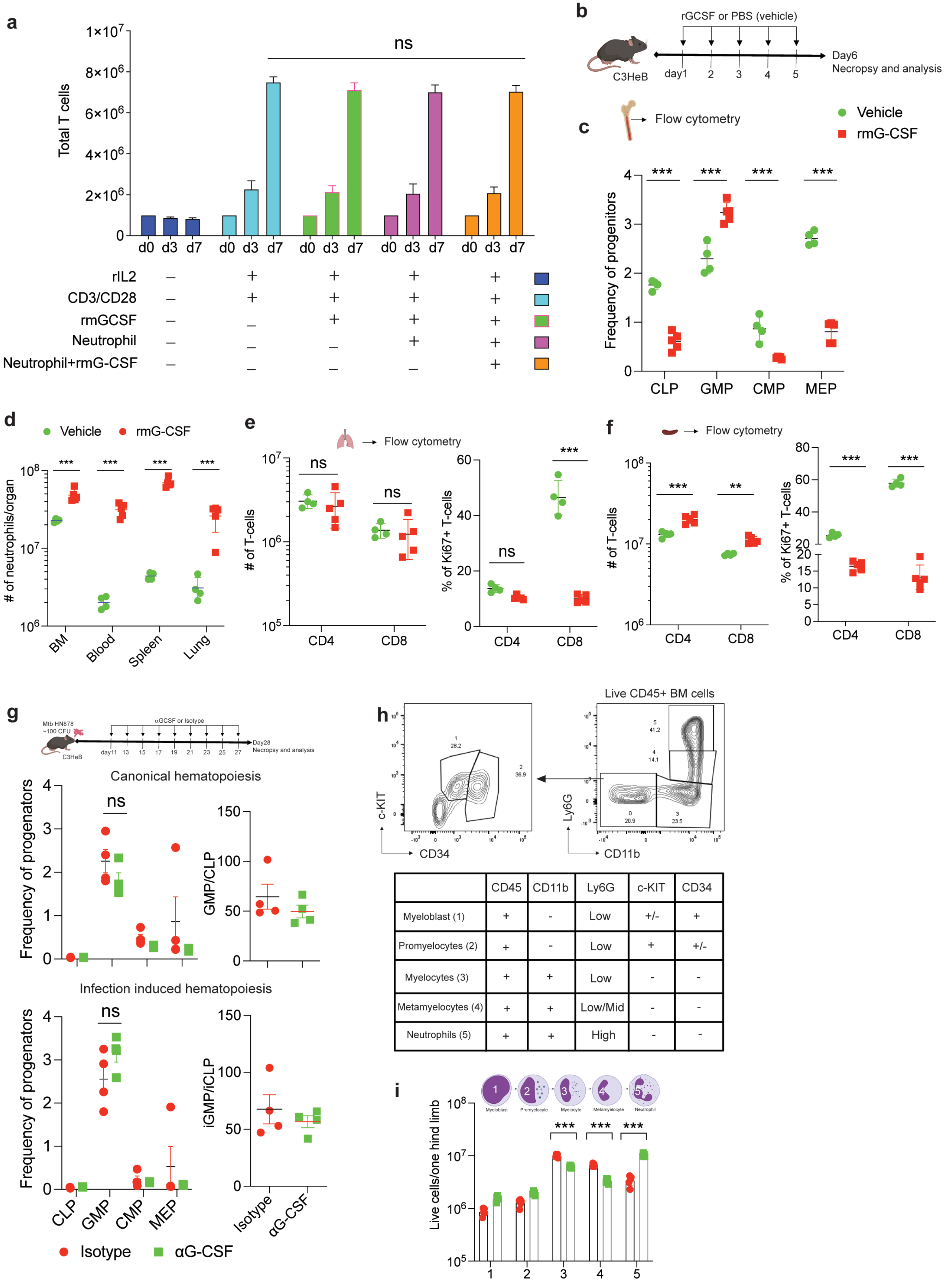


| | CD45 | CD11b | Ly6G | c-KIT | CD34 |
|---|---|---|---|---|---|
| Myeloblast (1) | + | - | Low | +/- | + |
| Promyelocytes (2) | + | - | Low | + | +/- |
| Myelocytes (3) | + | + | Low | - | - |
| Metamyelocytes (4) | + | + | Low/Mid | - | - |
| Neutrophils (5) | + | + | High | - | - |

### Extended Data Figure 7

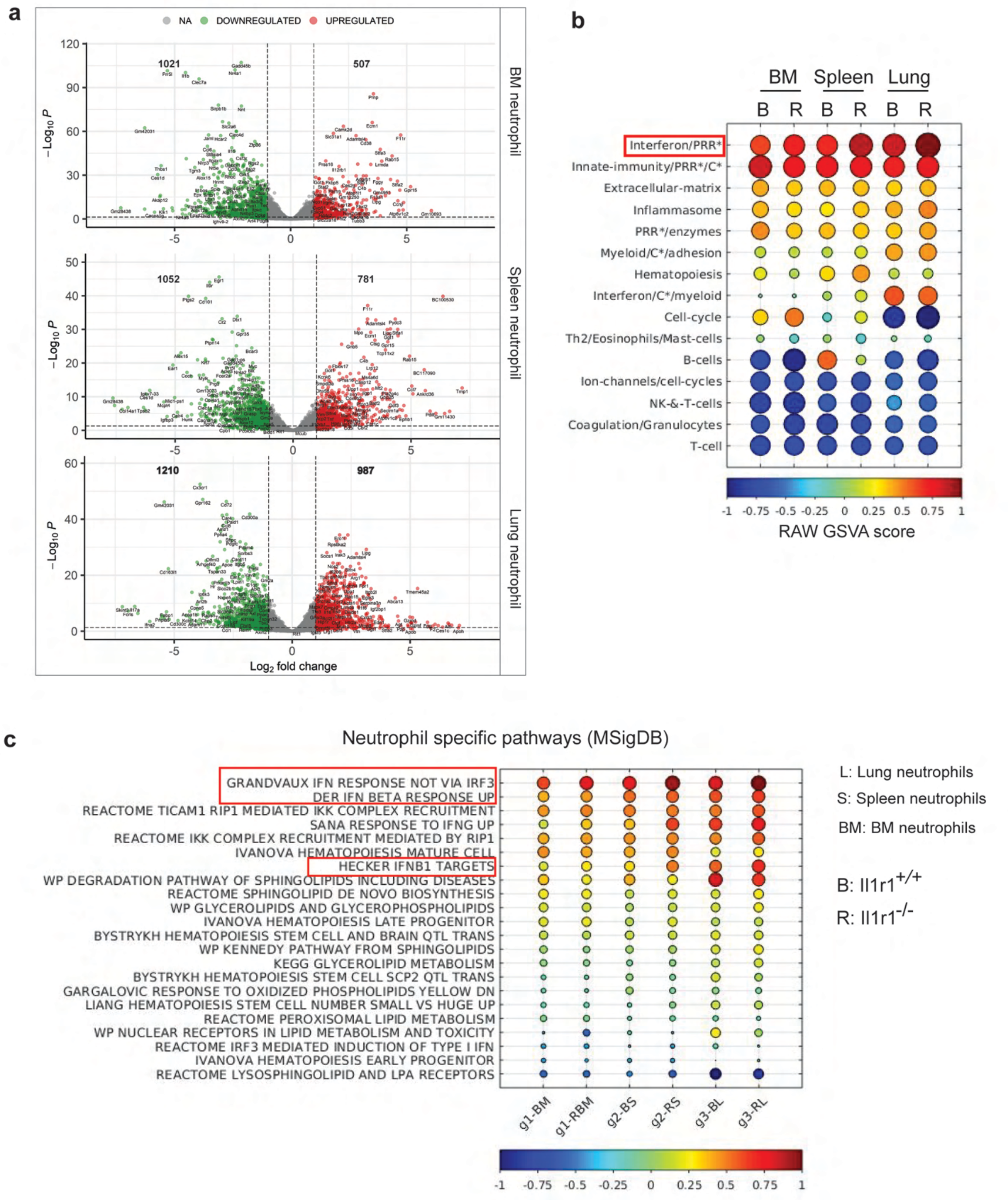

**Extended Data Fig. 8**

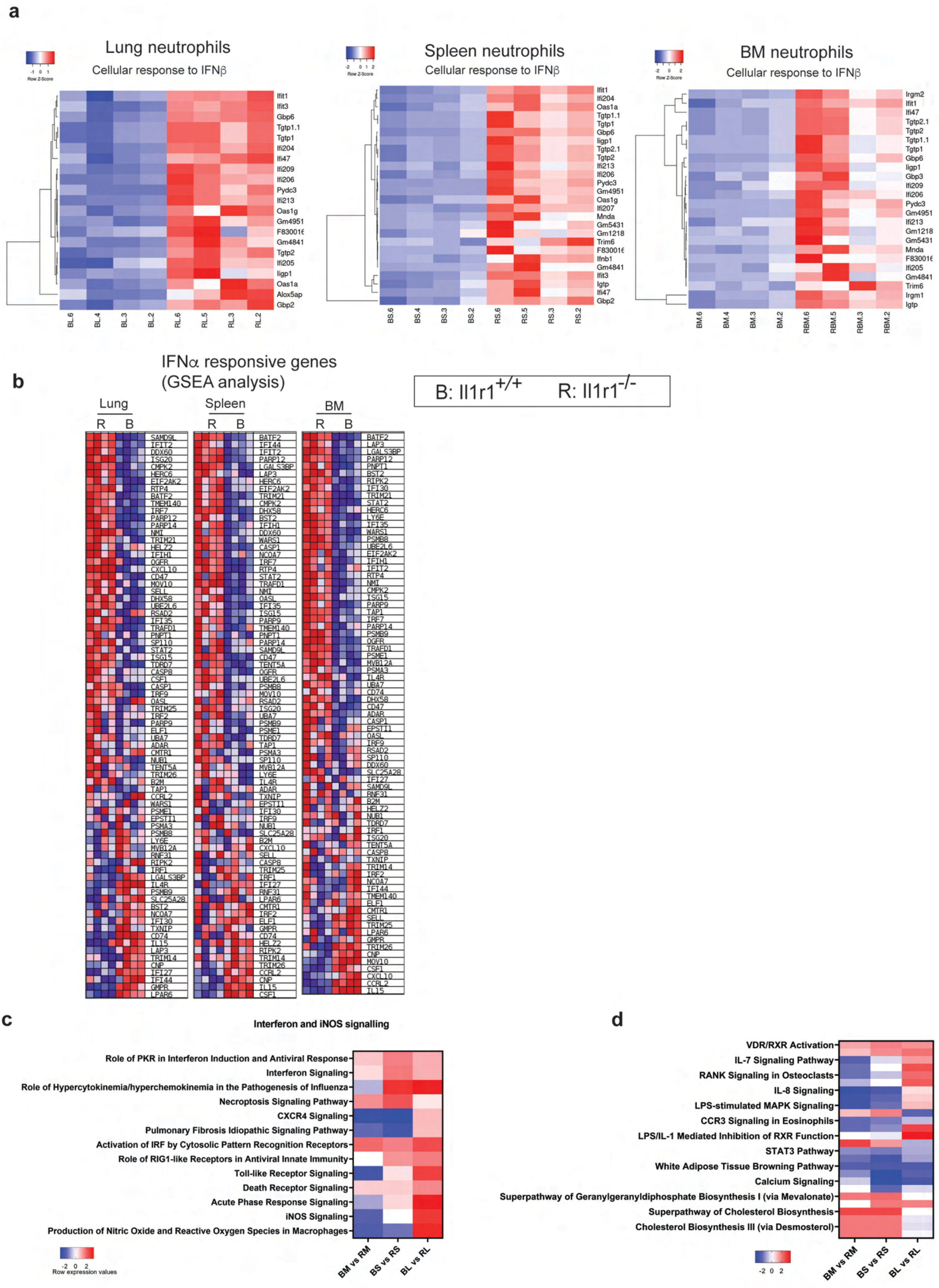

### Extended Data Figure 9

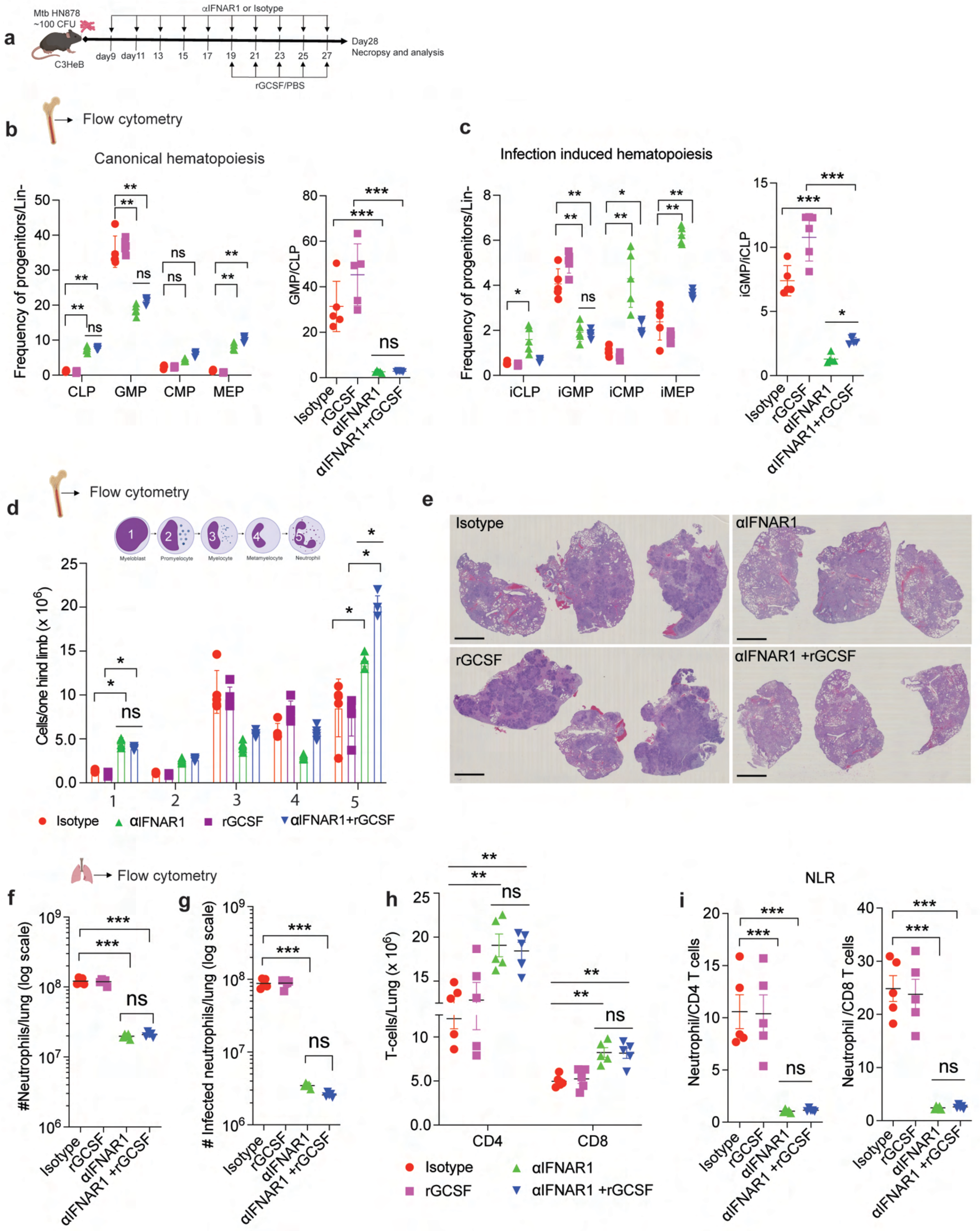

**Extended Data Figure 10**

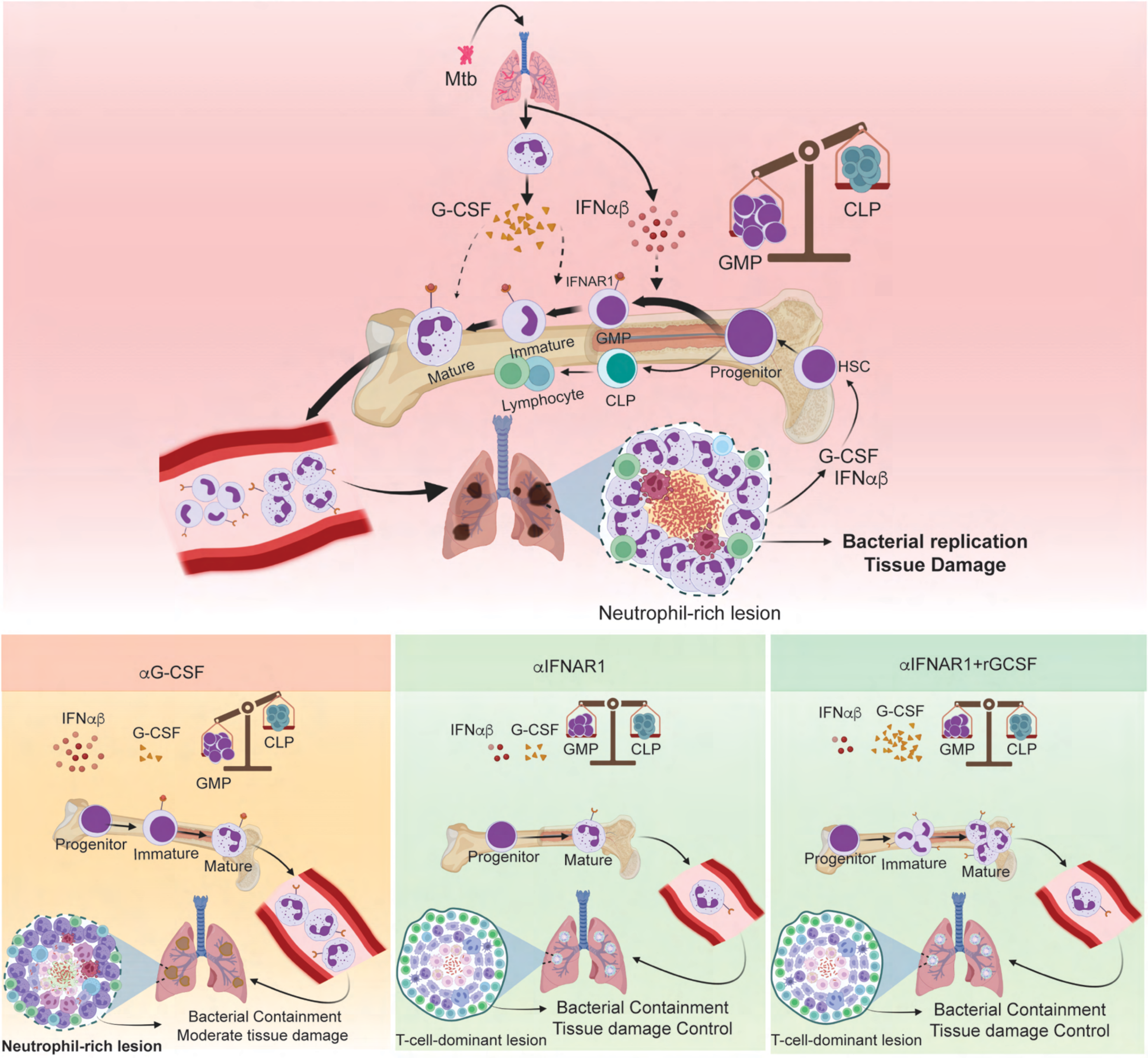
